# Repurposing the Memory-promoting Meclofenoxate Hydrochloride as a Treatment for Parkinson‘s Disease through Integrative Multi-omics analysis

**DOI:** 10.1101/2023.04.07.536024

**Authors:** Huasong Zhang, Cong Fan, Ling Li, Feiyi Liu, Shaoying Li, Linyun Ma, Yuanhao Yang, David N. Cooper, Yuedong Yang, Ronggui Hu, Huiying Zhao

## Abstract

Parkinson‘s disease (PD) is a devastating neurodegenerative disorder with growing prevalence worldwide and, as yet, no effective treatment. Drug repurposing promises to be invaluable for the identification of novel therapeutics for the treatment of PD due to the associated shortened drug development time, fewer safety concerns, and reduced costs. Here, we compiled gene expression data from 1,231 healthy human brains and 357 PD patients across ethnicities, brain regions, Braak stages, and disease status. By integrating them with multiple-source PD-associated genomic data, we found a conserved PD-associated gene co-expression module, and its alignment with the CMAP database successfully identified 15 drug candidates. Among these, we selected meclofenoxate hydrochloride (MH) and sodium phenylbutyrate (SP) for experimental validation because they are capable of passing through the blood brain barrier. In primary neurons, MH was found to prevent the neuronal death and synaptic damage associated with PD and to reverse the abnormal mitochondrial metabolism caused by PD. In hippocampal tissues, MH and SP were found to prevent the destruction of mitochondria, to reduce lipid peroxidation and to protect dopamine synthesis by PET-CT examination, malondialdehyde (MAD) testing and glutathione (GSH) testing, and immunohistochemical tests. Finally, MH was found to have the ability to improve gait behavior, and reduce anhedonic and depressive-like behaviors that are characteristics of PD mice. Taken together, our findings support the contention that MH may have the potential to ameliorate PD by improving mitochondrial metabolism and brain function.

## Introduction

Parkinson‘s disease (PD) is a chronic neurodegenerative movement disorder characterized by a large number of motor symptoms including resting tremor, rigidity and postural instability. The non-motor symptoms of PD include autonomic, psychiatric, sensory and cognitive impairments, as well as dementia[1]. As one of the most common neurodegenerative disorders, PD affects 2-3% of 65-year-olds and is responsible for more than 100,000 deaths worldwide each year[2]. However, our understanding of the etiology of PD remains incomplete. Thus, treatments for PD are still limited in their efficacy, e.g. dopamine replacement therapy, the most commonly used therapeutic strategy for PD, is capable of improving clinical symptoms but is unable to halt disease progression[3].

Thanks to decades of research, it is clear that PD exhibits considerable locus heterogeneity[4]; an increasing number of disease genes/pathogenic mutations are being identified in PD patients by means of whole genome sequencing (WGS) or whole exome sequencing (WES) studies[5], and both autosomal recessive and dominant forms have been described among the monogenic forms of PD. Such marked locus heterogeneity underlying PD not only represents a major obstacle in identifying common disease mechanisms, but restricts our options for appropriate therapeutic intervention. This notwithstanding, many mutations have been repeatedly identified among PD patients[6–8], thereby linking the familial and sporadic forms of PD mechanistically[9, 10]. Thus, mutations in the α-synuclein (*SNCA*) gene appear to be involved in both the familial and sporadic forms of PD; SNCA function/homeostasis is modulated by various contributory risk factors for PD, including oxidative stress, mitochondrial dysfunction, post-translational modifications and concentrations of fatty acids[11, 12]. It has therefore been reasoned that perturbation of the molecular networks involving multiple genes might commonly underlie the pathogenesis or progression of PD, and that the elucidation of these networks could facilitate the future development of therapeutic interventions. So far, the global effort toward this goal has led to the establishment of multiple valuable sources of information that has facilitated the compilation of such gene networks[13, 14]. Instead of focusing on single genes, considerable emphasis has been placed on utilizing data-driven frameworks at the system or network level to generate biologically/clinically meaningful gene modules comprising sets of functionally associated genes whose homeostasis may be altered by specific pathophysiological events[15–19]. The application of such an approach has led to the identification of network modules that are implicated in neurodevelopmental processes, metabolism and the immune system[20]. An analysis of GEO data of PD (*n* = 128) identified modules associated with RNA metabolism pathology as a potential cause of PD by sorting differentially active pathways between brain transcriptomics samples from PD patients and controls[21, 22]. However, these studies were generally based on patient gene expression data, and may have been biased due to insufficient numbers of samples and inter-patient heterogeneity.

One important application of the molecular network is in drug repurposing. Drug repurposing represents an attractive avenue in drug discovery due to its relatively low cost and fewer safety concerns. By definition, drug repurposing is designed to redirect new or additional indications for three kinds of therapeutic molecules i.e. drugs approved for a particular indication, drugs that have already been well-characterized during their clinical development and accompanied by thorough post-market surveillance data, and drugs which have undergone some clinical development but were subsequently abandoned [23, 24]. Often, biological networks combined with Genome-Wide Association Studies (GWAS) are the most commonly employed sources of information for drug repurposing, as GWAS studies are intended to impartially link controlled factor(s) to genetic or transcriptomic alterations in human subjects with no specific emphasis on a single gene or fixed set of genes[25, 26]. Thus, developing a method detecting the gene network perturbations caused by PD-associated variants through combining large-scale human genomic data, including functional interactions between genes from healthy human, is emerging as a useful approach to drug repurposing.

To test the effects of medicines for PD, an animal model is often used. The most common PD model involves a neurotoxin approach, such as the rotenone-induced PD model[27–29]. From 2000 onwards, researchers used rotenone to create a PD animal model[30], and it has proven to be very informative. It is thought to cause dopaminergic degeneration by inducing oxidative stress, as well as inducing *in vivo* aggregation of α-synuclein that is the major component of Lewy bodies[31]. Recently, Ahn et al. constructed a rotenone-induced PD mouse model in order to explore the role of δ-secretase in cleaving both α-Syn at N103 and Tau at N368[32]. Moreover, multiple studies have used the rotenone-induced mouse model in the study of PD-targeted medicines. For example, Liu et al. have investigated the protective effects of piperlongumine in rotenone-induced PD cell and mouse models[33]. Another study that used the rotenone-induced C57Bl/6J mouse model indicated the potential role of anle138b in the treatment of PD[34].

In an attempt to perform drug repurposing for PD, we have devised a computational architecture **i**ntegrating multiple-source **g**en**o**mic data with gene co-expression modu**l**es for **d**rug repurposing (iGOLD) (Fig. 1). It (https://github.com/fanc232CO/iGOLD_pipline) was applied to identify gene co-expression modules perturbed by PD-associated genes or SNPs, and to discover drugs that might be repurposed as PD therapeutics. The gene co-expression modules that were highly enriched in PD-associated genes and SNPs were further evaluated by module conservation analysis using seven gene expression datasets from multiple ethnicities, brain regions, tissues, Braak stages and PD disease status. The highly conserved module was used for drug discovery. Subsequent experiments with the rotenone-induced primary neuronal cells and the mouse model were conducted in order to validate the effectiveness of the drugs in promoting the survival of neurons, hippocampal functions and the modification of PD behavior. Finally, further experiments pertaining to mitochondrial functions and metabolic factors were conducted so as to reveal the specific mechanisms underlying the action of the drugs.

**Fig. 1.**
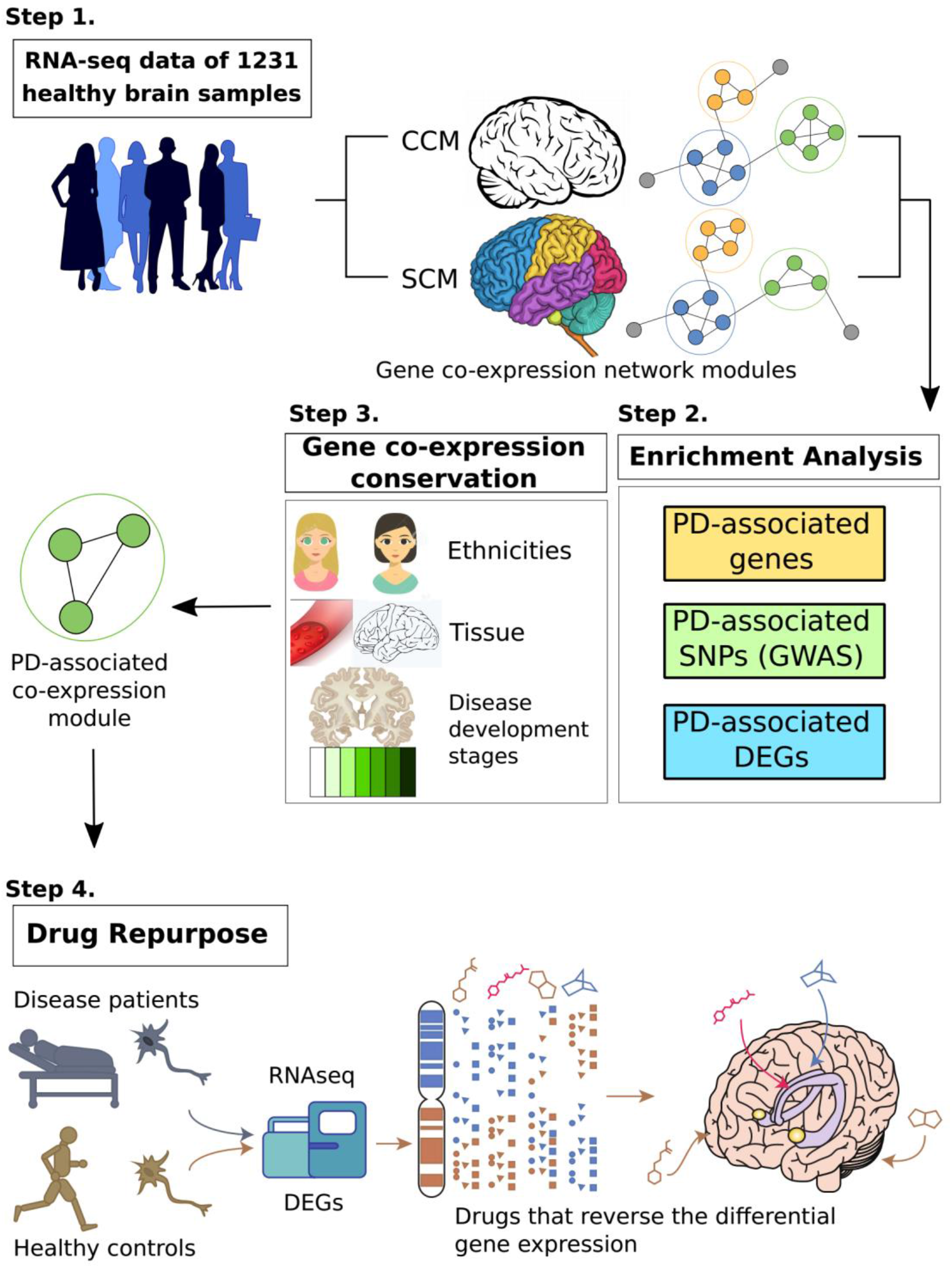
Schematics of iGOLD for drug repurposing. **Step 1:** Constructed gene co-expression modules using 1,231 healthy human samples from ten brain regions. CCM—concurrently co-expressed modules expressed in ten brain regions, SCM--brain region-**s**pecific **c**o-expressed modules. **Step 2:** From the constructed gene co-expression modules, we selected the PD-associated module through enrichment of PD-associated genes, enrichment of PD-associated SNPs (by Chi-square test and LDSC analysis, respectively) and fraction of PD-associated differential expression genes. DEGs—differential expressed genes. **Step 3:** Testing the conservation of the gene co-expression relationships in the validated modules across Ethnicities, Tissue and Disease development stages. **Step 4:** Inside the selected PD-associated module, the differential expressed genes in the PD patients were used for drug discovery with the Connectivity Map (CMAP). The enrichment of up-regulated and down-regulated genes by the drug-induced gene expression profiles were tested, and drugs that reverse the differential gene expression in PD were considered as the lead compound candidates.

## Results

### Overview of the study

Here, we firstly developed a computational architecture **i**ntegrating multiple-source **g**en**o**mic data with gene co-expression modu**l**es for **d**rug repurposing (iGOLD) for drug repurposing. The source code of iGOLD is available at (https://github.com/fanc232CO/iGOLD_pipline). As shown in Fig.1, it comprises by four main steps: (1) using gene expression data of 1,231 healthy human brain samples across 10 brain regions[35–37] to construct gene co-expression modules associated with normal brain functions; (2) identifying the co-expression modules enriched with disease-associated genes, SNPs and genes expressed significantly different in patients and controls; (3) examining the conservation of the selected co-expression modules in the gene expression data from brain tissues across different brain regions, disease status, and ethnicities, and in the gene expression data from blood and multiple other cell types; (4) aligning the highly conserved modules to the gene expression profiles perturbed by small molecular compounds in CMAP database [38, 39], and identifying the gene expression profiles enriched in genes from the conserved modules. The small molecular compound was considered as a drug candidate. The drug candidate was further validated by primary neuron and mouse models.

### The gene co-expression modules in hippocampi and substantia nigra as being associated with PD

Using the gene expression data from the healthy human brain, iGOLD constructed 19 **c**oncurrently **c**o-expressed **m**odules expressed in ten brain regions (CCM) (Table S1), and 68 modules (brain region-**s**pecific **c**o-expressed **m**odules, SCM) specifically expressed in one of the ten brain regions but not in the other nine brain regions (Fig. S1). The functional similarity of these modules was then evaluated by determining the number of overlapping genes between each pair of modules expressed in two brain regions. The width of the line in Fig. S1 represents the significance of the number of overlapping genes (Chi-square test) between one pair of modules from different brain regions compared to the modules from other brain regions (Table S2).

Among these modules, one CCM module, M3, and four SCM modules, BR7M4, BR9M3, BR6M3 and BR3MR, were suggested enriched (Bonferroni-corrected *P* < 0.05) in both PD-associated genes and PD-associated SNPs (Bonferroni-corrected *P_Chi-square_* < 0.05 and *P_sLDSC_*<0.05) (Fig. S2, Fig. S3, Table S1-S5). We further examined the enrichment of DEGs in these five modules (Fig. 2A). The DEGs were obtained from two GEO gene expression datasets[40], GPL96 and GPL97 (Table S6). The DEGs in GPL96 and GPL97 were respectively termed GPL96-DEGs and GPL97-DEGs. The proportions of GPL96-DEGs and GPL97-DEGs in BR7M4 are significantly higher than in other modules (Fig. 2A and Fig. S4), indicating that the BR7M4 module might best describe the gene expression profile characteristic of PD.

**Fig. 2.**
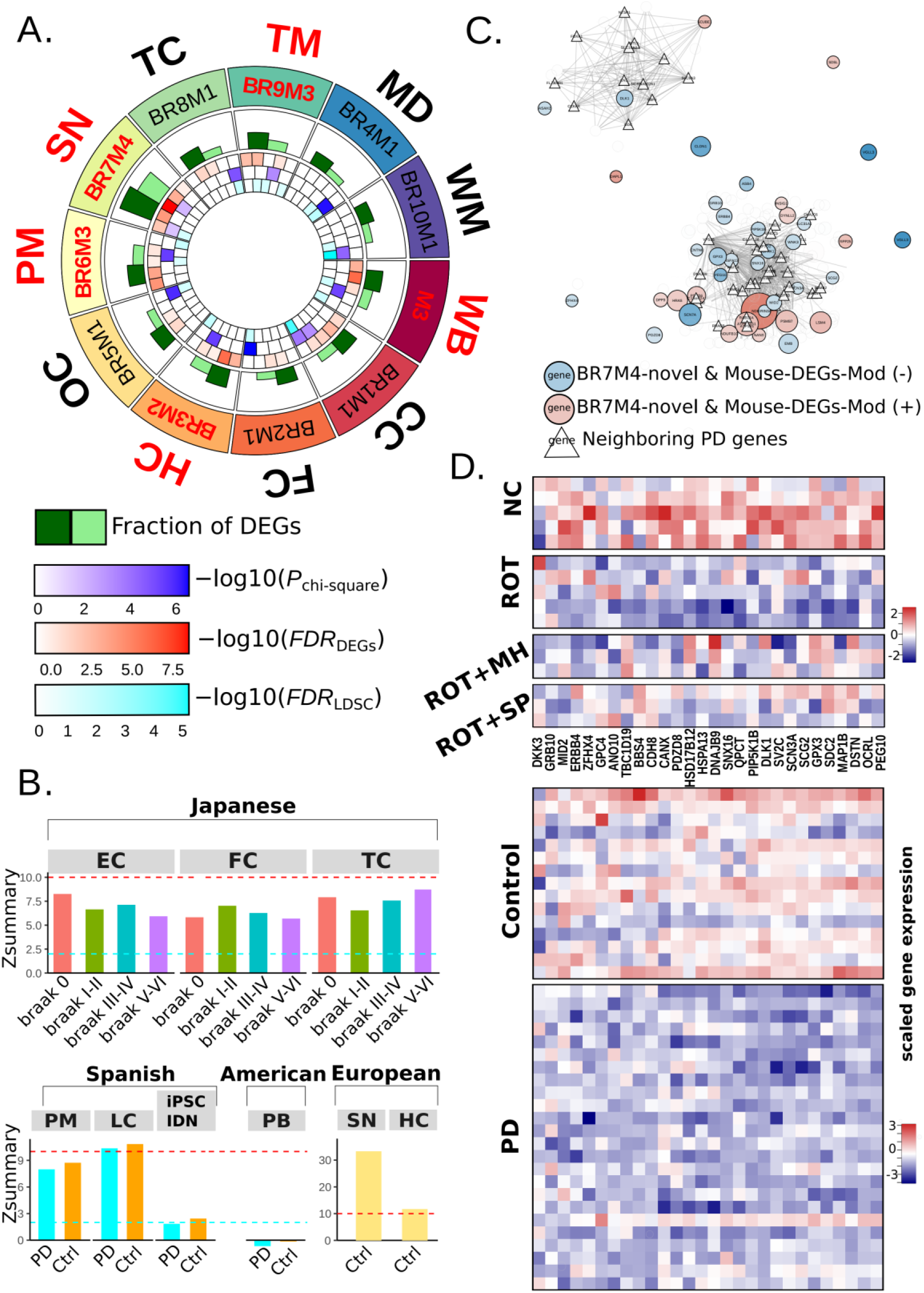
Detecting and validating gene co-expression modules associated with PD. (A) Enrichment of PD-associated genes and SNPs, and the proportion of differential expressed genes (DEGs) in co-expression modules that are most likely associated with PD compared to the other modules specifically expressed in the same brain region. From the outer ring to the inner, the circles sequentially represent the brain regions, the module names, the fraction of DEGs (dark green for GPL96 and light green for GPL97) in the co-expression modules, the enrichment of PD-associated genes by the co-expression modules, the enrichment of PD-associated SNPs by the co-expression modules tested by Chi-square analysis, and the heritability enrichment of PD-associated SNPs tested by sLDSC analysis. (B) Conservation of the BR7M4 genes in three brain regions of Japanese samples across different Braak stages including Braak0, BraakⅠ-Ⅱ, Braak Ⅲ-Ⅳ and BraakⅤ-Ⅵ. Conservation of BR7M4 genes expressed in brain regions, IPSC-induced dopaminergic neurons, and peripheral blood of PD patients and healthy controls. Gene co-expression conservation of BR7M4 module in the brain regions of hippocampus and substantia nigra, respectively. Red dashed line—high conservation Zsummary cutoff of 10. Cyan dashed line—medium conservation Zsummary cutoff of 2. (C) Interactions between BR7M4-novel genes and known PD-associated genes. BR7M4-novel genes overlapping with Mouse-DEGs are filled by red. Node size represents the significance of genes in RNA-seq analysis from the ROT group and NC group. The edge between the two genes represent their expression correlation less than 0.85 (scored by WGCNA), and genes linked by them are highlighted as triangles edged in black. (D) The gene expression profile of the NC group, ROT group, ROT+MH group and ROT+SP group. (E) The gene expression profile obtained by analyzing GPL96 data. Abbreviations: TC: Temporal Cortex, TM: Thalamus, WM: White Matter, CC: Cerebellar Cortex, FC: Frontal Cortex, HC: Hippocampus, MD: Medulla, OC: Occipital Cortex, PM: Putamen, SN: Substantia Nigra.

The associations between BR7M4 and PD were tested using seven unrelated publicly available brain expression datasets[35, 41–45] that together cover gene expression information across different ethnic backgrounds, brain regions, tissues and disease status (Table S7). As shown in Fig. 2B, module BR7M4 displays medium conservation in entorhinal cortex (EC), frontal cortex (FC) and temporal cortex (TC) at four Braak NFT stages (0, I–II, III–IV and V–VI) in Japanese samples. We examined the conservation of BR7M4 in putamen, locus coeruleus and IPSC-induced dopaminergic neurons using Spanish samples, and found that the BR7M4 module exhibits medium conservation in the putamen, high conservation in the locus coeruleus, and medium conservation in IPSC-induced dopaminergic neurons (Fig. 2B). When the BR7M4 module was tested in both hippocampus[35] and substantia nigra samples[35] from Europeans, it exhibited high conservation in both brain regions (Fig. 2B). By contrast, BR7M4 showed low conservation in the peripheral blood of American samples (Fig. 2B). Thus, unsurprisingly, the PD-associated functions of the BR7M4 module appear to be expressed through brain regions (e.g. substantia nigra and hippocampus) and dopaminergic neurons rather than through peripheral blood.

The module BR7M4 contained 399 genes whose interactions are shown in Fig. S5. The association of BR7M7 with PD was examined using hippocampal samples from mice since the expression of BR7M7 is highly conserved in the hippocampus (Fig. 2B, *Zsummary* = 10.53). We performed RNA sequencing (RNA-seq) on the hippocampi of ten into two groups of mice: DMSO (NC; *n* = 5) and ROT (rotenone-induced group; *n* = 5) (Table S8). The ROT-induced C57 L/J mouse model can recapitulate many features of human PD, including anatomical, neurochemical, behavioral and neuropathological features[32, 46, 47]. RNA-seq data analysis identified 2,195 genes (Mouse-DEGs) that were expressed significantly [adjusted *P* < 0.05, absolute value of fold-change (FC) greater than 2] differently between the NC and ROT groups. Among the Mouse-DEGs, 50 genes were present in the BR7M4 module (termed Mouse-DEGs-Mod), which is significantly (single-tailed binominal test *P* = 0.016) more than the genes that were not expressed significantly differently between the NC and ROT groups (termed Mouse-non-DEGs) (Fig. S5). Moreover, 39 of the Mouse-DEGs-Mod genes were not PD-associated genes. Nevertheless, the expression levels of these 39 genes were found to be closely (WGCNA TOM similarity > 0.15) correlated with those of 41 PD-associated genes in BR7M4 (Fig. 2C). Thus, the co-expression module, BR7M4 that is strongly associated with PD.

### Meclofenoxate hydrochloride and sodium phenylbutyrate restore the normal expression levels of PD-associated genes via different mechanisms

From BR7M4, we extracted DEGs for the discovery of PD candidate therapeutics (Table S9), from which two drugs, sodium phenylbutyrate (SP) (connectivity score-0.963 and ranked in top 0.05% of 6100 drugs) and meclofenoxate hydrochloride (MH) (connectivity score of-0.814 and ranked in top 0.2% of 6100 drugs), were considered for further validation since they were not only top-ranking candidates but were also able to pass through the blood brain barrier.

To assess the impact of MH or SP on PD-associated gene expression, we performed RNA-sequencing on the hippocampi of mice from the NC, ROT, ROT+SP, ROT+MH, SP and MH groups (Table S8). We found 91 genes to be expressed significantly (Bonferroni-corrected *P* < 0.05 and |log(FC)| > 2) differently between the ROT group and the NC group as well as between the ROT+SP group and the ROT group. These genes were termed the SP-ROT set. Meanwhile, we found 666 genes that were expressed significantly (Bonferroni-corrected *P* < 0.05 and |log(FC)| > 2) differently between the ROT group and the NC group as well as between the ROT+MH group and the ROT group. These genes were termed the MH-ROT set. Among them, 28 were in the BR7M4 module and displayed the same direction of regulation as the GPL96 dataset. The expression of these genes in the ROT group, the NC group, the ROT+MH group and the ROT+SP group are shown in Fig. 2D. The expression of these genes in the GPL96 dataset is shown in Fig. 2E. The expression profile of these gene in controls in GPL96 is similar to that of the NC group, whereas the gene expression profile of the PD individuals in GPL96 is similar to that of the ROT group (Fig. 2D, Fig. 2E). Thus, after MH or SP treatment, gene expression in the ROT group was restored such that it approximated to the characteristics of the NC group, suggesting a specific effect of MH or SP in remodeling the expression pattern of PD-associated genes.

Of the genes in the MH-ROT set, 129 genes were not in the SP-ROT set whilst 74 genes from the SP-ROT set were not in the MH-ROT set, which were then termed the Uni-MH-ROT set and Uni-SP-ROT set, respectively. A STRING analysis was performed to detect the networks of protein-protein interactions (PPIs) in the Uni-MH-ROT set and the Uni-SP-ROT set, respectively. The PPIs of the Uni-MH-ROT genes were mainly enriched in synapse-related functions (Fig. S6A) whereas the PPIs of the Uni-SP-ROT group were enriched in mitochondrial electron transport and mitochondrial respiratory chain complex I assembly functions (Fig. S6B). Thus, the effect of MH on gene transcription is potentially distinguishable from that of SP in terms of its modulatory effect on genes with synapse-related functions in murine hippocampus. The difference between MH and SP was further examined by DStruBTarget[48] to predict those proteins that could directly bind to MH and SP (Supplementary Material and Table S10). DStruBTarget indicated that the top 10 predicted proteins binding to MH were enriched in neuroactive ligand-receptor interactions and neurotransmitter receptor activity functions (*P* =1.3× 10^-7^), whereas the top 10 DStruBTarget predicted proteins binding with SP were enriched in inflammation-related functions (Table S11). Thus, MH and SP may bind to different targets for restoring the normal expression levels of PD-associated genes in the hippocampi of mice.

### Both SP and MH protect neurons against ROT-induced neurodegeneration

The neuronal nuclear protein (NeuN) is often used as a positive marker for the functional state of postmitotic neurons. Thus, the NeuN-positive rate of neurons is usually used to assess neurodegeneration[49–51]. Here, immunohistochemical (IHC) staining with anti-NeuN was performed on the dentate gyrus (DG), dentate gyrus2 (DG2) and cornu ammonis (CA1) of the hippocampus from six groups of mice (NC, ROT, SP, ROT+SP, MH and ROT+MH), with four mice in each group. As shown in Fig. 3A and B, the average relative numbers of NeuN-negative cells [quantified by ImageJ[52]] (29.52%) increased by 27.89% in the ROT group as compared to those of the NC group (1.63%) (*P* = 2.7× 10^-3^). In the SP+ROT and MH+ROT groups, the average relative numbers of NeuN-negative cells (3.08% for SP treatment and 2.38% for MH treatment) were respectively reduced by 26.44% and 27.14% (*P* = 1.9× 10^-3^, and *P* = 1.3× 10^-2^, respectively) compared to the ROT group (Fig. 3A and 3B). The average relative numbers of NeuN-negative cells in the DG2 structure of the hippocampus in the ROT group (30.56%) increased by 26.69% compared to the NC group (3.88%) (*P* = 1.5× 10^-2^). In the SP+ROT and MH+ROT groups, the average relative number (5.18% and 4.55%, respectively) of NeuN-negative cells in DG2 was reduced by 25.38% (*P* = 2.3 × 10 ^-2^) and 26.01% (*P* =3.6 × 10 ^-2^), respectively, compared to the ROT group (Fig. 3A and 3C). In the CA1 substructure of the hippocampus, the average relative number (54.73%) of NeuN-negative cells in the ROT group increased by 50.87% compared to the NC group (3.86%) with *P* = 1.7× 10^-2^. The average relative numbers of NeuN-negative cells of ROT+SP (3.00%) and ROT+MH (3.08%) in the CA1 substructure of the hippocampus were reduced by 51.73% and 51.65%, respectively (*P* =1.8× 10^-2^ and *P* =3.5× 10^-2^) compared with the ROT group (Fig. 3A, and 3D). Thus, MH and SP treatments reduce the numbers of NeuN-negative cells in different parts of the hippocampus.

**Fig. 3.**
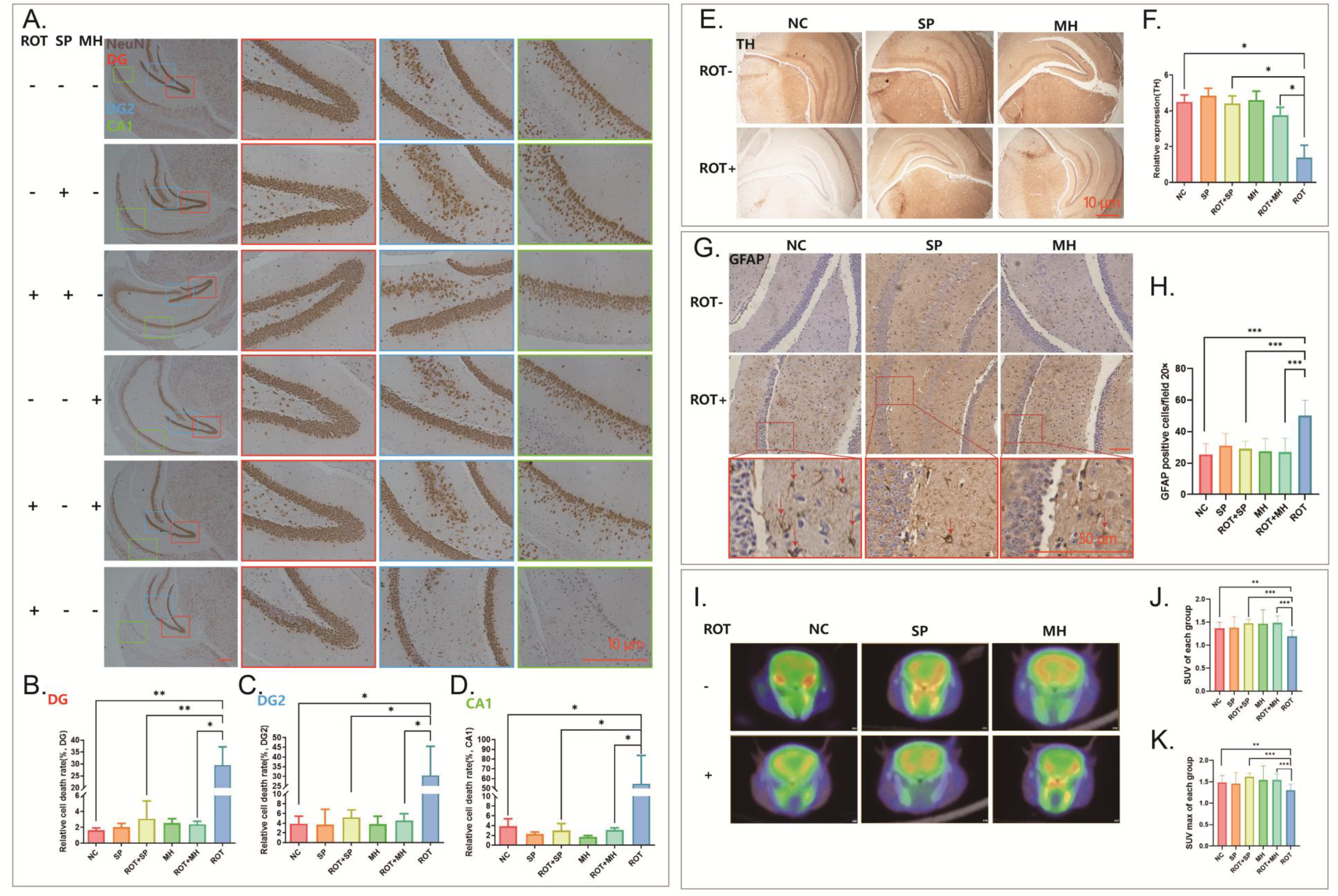
MH and SP can protect neurons against PD-related neurodegeneration. (A) Immunohistochemical representation of NeuN in the DG, DG2 and CA1. Magnification 20×. Scale bar = 10μm. (B) The relative number of NeuN-negative cells in the DG structure of the hippocampus in different groups. (C) The relative number of NeuN-negative cells in the DG2 structure of the hippocampus in different groups. (D) The relative number of NeuN-negative cells in the CA1 structure of the hippocampus in different groups. (E) Immunohistochemical representation of TH in the hippocampus of the experimental mice (N ≥ 3 mice/group). TH expression was reduced in the ROT-induced mice, and increased in the SP-treated and MH-treated mice as compared with the NC. Magnification 4×. Scale bar = 10μm. (F) The relative expression of TH in the ROT group was significantly lower than that in the NC group. SP treatment and MH treatment can increase the relative expression of TH in the ROT-induced group. (G) GFAP in the hippocampus of each group. The number of GFAP-labelled astrocytes was significantly increased in the ROT group as compared to the NC. The number of GFAP-labelled astrocytes was significantly reduced in the ROT+SP-treated mice and ROT+MH-treated mice as compared to the ROT group. Magnification 10×. Scale bar = 50μm. (H) The relative number of GFAP-labelled astrocytes in the hippocampus structure in different groups. (I) Fluorescence image of glucose metabolism capacity shown by PET of mouse brain. (J) The average change of SUVs in each group. (K) The maximum change of SUVs in each group. SP and MH prevented ROT-induced neurodegeneration. Abbreviations: [18F]-FDG, ^18^F-fluorodeoxyglucose; α-SYN: α-synuclein group; eGFP, enhanced green fluorescent protein; PET, positron emission tomography; A, anterior; P, posterior; L, left; R, right. \**P* < 0.05, \*\**P* < 0.01, \*\*\**P* < 0.001.

In neurons, the expression level of tyrosine hydroxylase (TH) is generally considered to be an indicator of the ability of cells to produce dopamine. IHC analyses were therefore performed in order to examine the relative expression levels of TH in the hippocampal tissues of the mice. As shown in Fig. 3E and 3F, the IHC staining signal of TH in the SP+ROT and MH+ROT groups were more intense than in the ROT group. The expression level of TH in the ROT group (relative expression of TH=1.40) was lower than that of the NC group (relative expression of TH=4.49) (*P* = 1.8× 10^-2^). In the SP+ROT group, the expression of TH was 4.41, which is over 3-fold higher than in the ROT group (*P* =1.4× 10^-2^). Similarly, the expression level of TH in the ROT+MH group was 3.75, more than two-fold higher than in the ROT group (*P* = 1.6× 10^-2^) (Fig. 3F). Thus, both SP and MH treatment improved the ability of neurons to produce dopamine.

Subsequently, IHC staining with anti-GFAP (glial fibrillary acidic protein) was also performed to visualize the intermediate filament (IF) protein expressed in numerous cell types of the central nervous system (CNS) including astrocytes and ependymal cells, with the number of GFAP-positive (GFAP^+^) cells serving as an indicator for the activation of the neuroinflammatory pathway in the murine hippocampus (Fig. 3G). As shown in Fig. 3G, the proportion of GFAP^+^ cells was markedly increased in the hippocampus of the ROT group (50.17±9.88) compared to that in the NC group (25.50±6.80)(*P* =3.83 × 10 ^-7^). By contrast, the numbers of GFAP^+^ cells in the ROT+SP and ROT+MH groups were significantly lower than that of the ROT group [SP: 29.08 ± 4.81 (*P* =1.16× 10^-6^) and MH: 26.89 ± 8.87 (*P* =2.23× 10^-5^)]. However, little or no difference was observed in terms of the numbers of GFAP^+^ cells between the ROT+SP and ROT+MH groups (Fig. 3H). Similar results were obtained for the interleukin 1 complex (IL-1), another proinflammatory cytokine and GFAP, in the murine striatum (Fig. S7). Taken together, it is clear that both MH and SP repress ROT-induced neuroinflammation in the hippocampus, suggesting an anti-inflammatory effect of these drugs.

### MH and SP both up-regulate glucose metabolism in the brains of ROT-induced PD mice

To measure glucose metabolism in mouse brains, mice from both the NC and ROT groups (Table S8) were subjected to neuro-imaging through [18F]-fluorodeoxyglucose positron emission tomography (^18^F-FDG PET) (Supplementary Materials). The cross-sectional small animal PET images of mice from the six groups are presented in Fig. 3I and are quantified in Fig. 3J. In Fig. 3J, the average and maximum standardized uptake value (SUV) of the ROT group are 1.19 and 1.30 which were decreased by 0.18 (*P* =2.3× 10^-2^) and 0.18 (*P* = 3.2× 10^-2^), respectively compared to the NC group (average SUV = 1.37 and maximum SUV = 1.48) (Fig. 3K), indicating higher intensity of [18F]-FDG uptake in the ROT-induced group than in the control group (NC). These data clearly indicate that ROT treatment markedly down-regulated glucose metabolism of the murine neurons. For the mice in the SP+ROT group, the average SUV was 1.48 (Fig. 3J), whilst the maximum SUV value was 1.62 (Fig. 3K), which were decreased by 0.28 (24.37%, *P* = 2.67 × 10^-6^) and 0.32 (24.62%, *P* = 1.67 × 10^-6^), compared with the ROT group. For the mice in the ROT+MH group, the average SUV value was 1.48, with a maximum SUV value of 1.54, 0.32 (24.62%) and 0.24 (18.46%) higher than for the ROT group (*P* = 5.04 × 10^-5^ and (*P* = 5.29 × 10^-4^), respectively (Fig. 3J and 3K). Thus, treatment with either SP or MH appears to significantly promote glucose metabolism in neuronal cells in ROT-induced PD mice.

### Validating the action of SP and MH in preventing ROT-induced cell damage and preserving neuronal cell morphology

We then tested the potential effects of SP and MH on the survival and morphology of cells in primary neuron culture. The proportions of viable cells as well as the relative volume of cell bodies were calculated by ImageJ[52]. The volumes of the neuronal cell bodies appeared to shrink to 25.92% of the NC in the presence of ROT (Fig. 4Aa and Fig. 4Ab). After SP and MH treatment, the volumes of the neuronal cells increased by 4.78 (*P* =7.9× 10^-3^) and 6.59 (*P* =5.1× 10^-3^), respectively, compared to the ROT group (Fig. 4Ab). The average number of surviving cells in the ROT-induced group (9.14) was significantly lower than in the NC group (154.43) (*P* = 6.2 × 10^-4^) (Fig. 4B). Remarkably, the treatment of the ROT-induced cells with either SP or MH increased cell survival from 9.14 to 63.71 or 47.43, respectively (Fig. 4B), strongly indicating that SP and MH has the potential to protect neurons from ROT-induced damage.

**Fig. 4.**
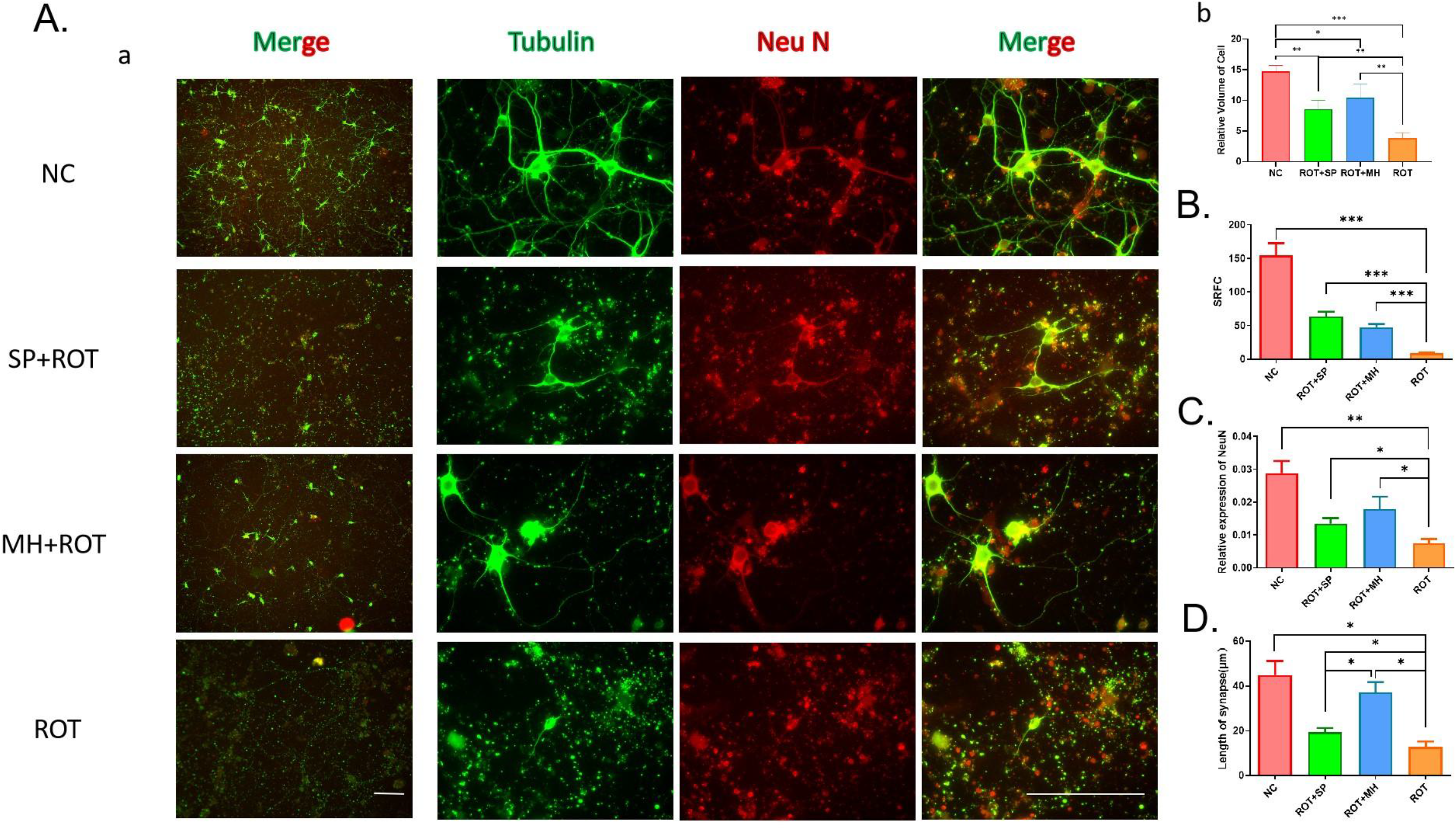
SP and MH prevent ROT-induced neuronal cell damage and protect neuronal cell morphology. (A) a: Representative scans from immunocytochemical preparations acquired with 4× and 100× objective lenses. SP protects NeuN+ and mouse neurons against the deleterious effects of ROT with an ensuing increase in NeuN and tubulin expression in PD-containing neurons. Insets correspond to high magnification images. Data are representative of 20–40 neurons per group obtained from seven independent cultures. b: The relative volume of cells in the ROT group was significantly smaller than that in the other three groups. (B) The number of tubulin+ cells with cell bodies and synapses in more than three visual fields of each group. SP and MH effectively prevent cell death caused by ROT-induced cytotoxicity and increase the number of viable cells in each field. (C) The relative expression of fluorescence of the ROT group was significantly lower than that of the NC group. The relative expression of fluorescence increased after SP and MH treatments (ROT+SP and ROT+MH groups). (D) The length of the nerve synapse of the ROT group was significantly lower than that of the NC group, and the length of the nerve synapses increased after SP and MH treatments (ROT+SP and ROT+MH groups). Scale bar: 20μm in 4× and 500μm in 100×. On average, 25-30 neurons per condition were tested from three independent cultures. \*\*\**P* < 0.001, \*\**P* < 0.01 and \**P* < 0.05.

Furthermore, the relative expression of tubulin fluorescence (fluorescence/area) in the ROT-induced group was 0.007, significantly lower than that in the NC group (*P* = 2.0 × 10 ^-2^), which increased to 0.013 and 0.018 after SP and MH treatment, respectively, significantly higher than in the ROT-induced group (*P* = 1.8× 10^-2^ and 1.2× 10^-2^, respectively) (Fig. 4C). Moreover, we found that the average lengths of the nerve synapses in the ROT-induced group were approximately 12.75μm, significantly shorter than that in the NC group (∼44.98μm) (*P* = 1.3× 10^-2^). After SP and MH treatment, the lengths of the nerve synapses increased to 20.19μm (*P* = 2.6 × 10^-2^) and 37.06μm (*P* = 2.3× 10^-2^), respectively, significantly higher than for the ROT-induced groups. Interestingly, the average length of the synapses in neurons treated with MH was longer than that treated by SP (*P* =2.1× 10^-2^) (Fig. 4D).

### MH and SP improved gait, anhedonia and the depression-like behaviors of PD mice

To further assess the potential of MH and SP as drug candidates to treat PD, we evaluated to what extent MH or SP might impact PD-like behavior traits in the ROT-induced mouse model, where ROT is known to cause a gait abnormality. The mice from each of the abovementioned six groups were subjected to a gait behavior test (for detailed description on the test, see Supplementary Material). As shown in Fig. 5A, the footprints of the NC group (red) proceeded in a straight line, whereas the footprints of the ROT group (orange) were not straight, indicating an unstable gait. The gait behavior of the SP-(yellow) or MH-treated (cyan) groups was also assessed in comparison with that of the NC or ROT group. In the ROT+SP group (green), the footprints were almost straight although the walking directions of the front and rear feet on the same side were not exactly parallel to each other. By comparison, for the mice in the ROT+MH group (blue), the front and rear feet on the same side were precisely in parallel, with the lines of the footprints straighter than those of the mice in the ROT+SP group (Fig. 5A).

**Fig. 5.**
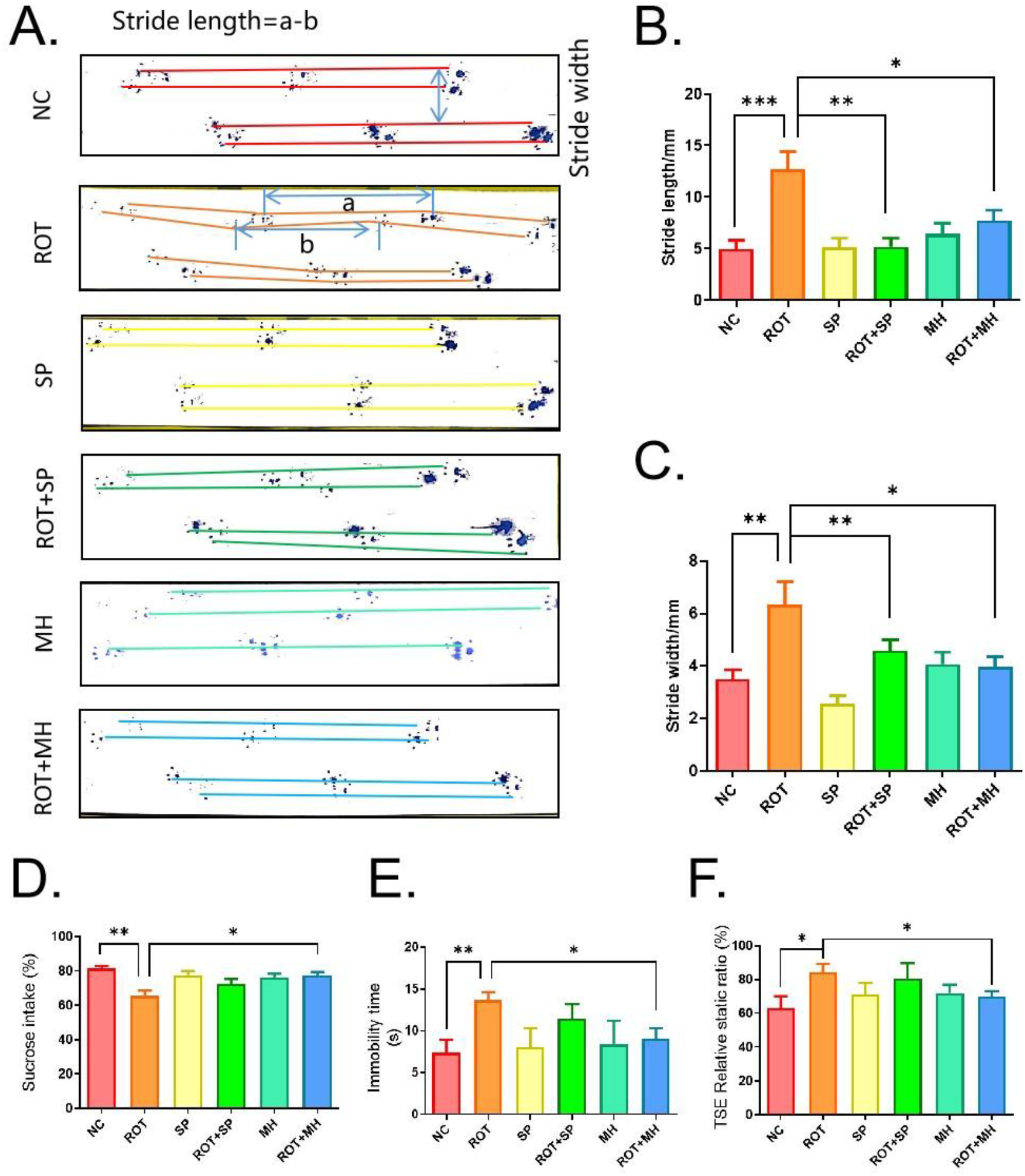
MH can ameliorate PD-related behaviors. (A) The influence of SP and MH on the behavior of the PD animal model constructed by ROT (footprinting test). PD behavioral changes were mainly disclosed as gait changes, whilst the repetition rate of footprints decreased. (B) Stride length of mice. The stride length of PD mice treated with ROT+SP and ROT+MH was significantly less than that of PD mice. (C) Stride width of mice. The stride width of mice treated with ROT+SP and ROT+MH was significantly less than that of PD mice, respectively. (D) Effects of SP and MH pretreatment on ROT-induced anhedonic behavior evaluated by means of the sucrose preference test (SPT). (E) Effects of SP and MH pretreatment on ROT-induced depressive-like behavior of mice evaluated by the forced swim test (FST). (F) Effects of SP and MH pretreatment on ROT-induced depressive-like behavior of mice evaluated by the tail suspension experiment (TSE). \**P* < 0.05, \*\**P* < 0.01, \*\*\**P* < 0.001.

To quantify the gait behavior of the mice, we measured the stride lengths of the mice. As shown in Fig. 5B, the stride lengths of the NC, SP and MH groups were 5.00mm, 5.18mm and 6.46mm, respectively, with no statistically significant difference evident between them (*P* > 0.05). The average stride length of the ROT group was however 12.67mm, 2.5 times longer than that of the mice in the NC group (*P* = 2.0× 10^-3^). SP treatment served to completely reverse the ROT-induced increase in stride length, as the average stride length of the ROT+SP group was 5.22mm, almost 2.5 times shorter than that of the ROT group (*P* =1.0× 10^-4^). For the ROT+MH group, the average stride length was 7.73mm, also significantly smaller than that of the ROT group (*P* =1.0× 10^-4^) (Fig. 5B). Turning to the stride width, those of the NC, SP and MH groups were 3.49mm, 2.55mm and 4.06mm, again not statistically different from each other (*P*>0.05) (Fig. 5C). Whilst the average stride width of the ROT group was 6.35mm, significantly greater than for the mice in the NC group (*P* =1.0× 10^-3^), the average stride widths measured for the SP+ROT and MH+ROT groups were 4.59mm (*P* =1.4× 10^-3^) and 3.97mm (*P* =1.7× 10^-2^), respectively, suggesting a narrowing effect of SP and MH on the stride width of the animals (Fig. 5C). Taken together, these data strongly indicate that the administration of SP or MH can efficiently restore the ROT-induced changes in mouse gait behavior, suggesting that these drugs might offer some hope of alleviating the gait abnormality typically observed in PD patients.

Clinically, anhedonia (lack of interest) and depression are commonly noted in PD patients. Whilst a reduction in the sucrose preference ratio in an experimental group compared to controls is held to be indicative of anhedonia in animals[53], the forced swimming test (FST) and the tail suspension test (TST) have been devised to assay depression-like behavior in the preclinical mouse model for PD[54, 55]. The sucrose preference test was performed to measure the percentage of sugar water intake in 24 hours for the mouse groups (Supplementary Material). The sucrose preference of the ROT group (65.37%) was approximately 25% lower than that of the NC group (81.37%) (*P* =2.8× 10^-2^). Interestingly, the sucrose preference of the ROT+MH group (77.54%) increased by approximately 20% compared to the ROT group (*P* =2.8× 10^-2^), whilst the sucrose preference of the ROT+SP group increased to 72.36%, but this increase was subtler and did not attain statistical significance (*P* =1.2× 10^-1^) (Fig. 5D).

In the forced swimming test (FST), the ―immobility‖ time indicates the states the experimental animals eventually adopt to avoid the stressor (water, in this case), which could be quantified to indicate depression-like behavior of mice[54]. As shown in Fig. 5E, the immobility time for the ROT group (13.86s) was longer than for the NC group (7.38s) (*P* =8.0× 10^-3^). The immobility time of the ROT+MH group was significantly lower (9.07s) than for the ROT group (*P* =1.5× 10^-2^). However, the average immobility time (11.42s) of the ROT+SP group was comparable to that of the mice that received only ROT (Fig. 5E).

The tail suspension experiment (TSE) was quantified using a ratio of static time to moving time. As shown in Fig. 5F, the ratio of static time to moving time in the ROT group (84.49%) was approximately 20% greater than in the NC group (63.01%) (*P* =3.3× 10^-2^). Remarkably, the ratio of static time to moving time in the ROT+MH group (70.00%) was significantly lower than in the ROT group (*P* =3.2 × 10 ^-2^), whereas the average ratio of static time to moving time (80.52%) in the ROT+SP group scarcely changed from that of the ROT group (Fig. 5F).

Taken together, we conclude that MH can significantly alleviate both anhedonia and depression-like behavior in the ROT-induced mouse model for PD, with considerably greater efficiency than SP.

### Exploring the mechanisms by which MH protects mitochondrial function, postsynaptic density and synaptic functions in neuronal cultures and mouse brain

Having found that the impact of MH on the behavior of PD mice was significantly greater than that of SP (Fig. 5A-F), we next directed our efforts toward elucidating the underlying mechanisms by which MH treatment might be efficacious. We therefore sought to ascertain the impact of MH on mitochondrial homeostasis and function in the hippocampus of the ROT-induced PD mouse model. The mitochondria presented themselves as continuous filaments in the hippocampal neurons of the NC group whilst they showed more obvious fragmentation after treatment with ROT (Fig. 6Ac). Additionally, ROT treatment appeared to lead to a concentration of clumped mitochondria in the cell bodies of the neurons that were broken and lumpy in shape (Fig. 6A panel a-c). Remarkably, compared to the disrupted mitochondria in the neurons from the ROT group, the ROT+MH group appeared to have significantly restored the mitochondrial morphology in the neurons, with most of them being maintained in the soma, and continuous in shape (Fig. 6Ae,f). Interestingly, when the neuronal cells were treated with MH alone, mitochondria were observed with many more synapses in the neurites than in the NC group (Fig. 6Ad).

**Fig. 6.**
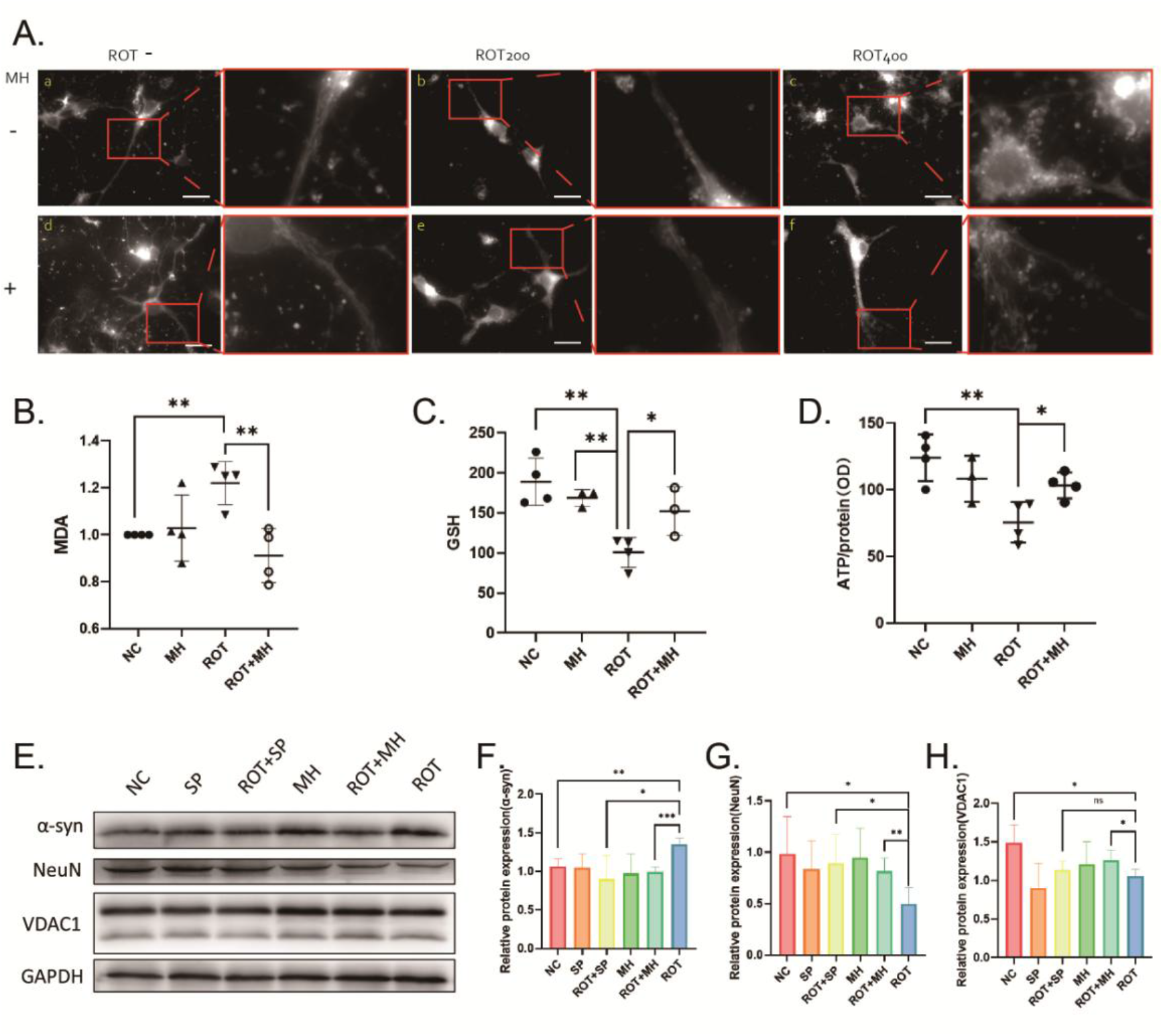
MH can reduce mitochondrial damage in neurons. (A) Protective effects of MH on mitochondrial morphology in a ROT-induced PD cell model. a: Mitochondrial morphology of control neurons. b: Mitochondrial morphology in a ROT-induced PD cell model with 200μM ROT. c: Mitochondrial morphology in a ROT-induced PD cell model with 400μM ROT. d: Mitochondrial morphology of neuronal cells treated with MH alone. e: Mitochondrial morphology of the PD neuron model induced by 200μM ROT and treated with MH. f: Mitochondrial morphology of the PD neuron model induced by 400μM ROT and treated with MH. (B)-(E): MH protecting mitochondrial function in neurons. Effects of MH pretreatment on ROT-induced oxido-nitrosative stress assessed by (B) malondialdehyde (MDA) level, (C) reduced glutathione (GSH) level, and (D) ATP levels of the hippocampus of the mice in each group. (E)-(H): According to the results of mitochondrial dysfunction, we performed western blotting for mitochondrial-related proteins. The western blot (E) was quantified for α-syn(F) and NeuN(G) in the murine hippocampi. The bands were quantified using Sigma Gel software, and the differences are represented by a histogram. GAPDH was used as a loading control. The results showed that both MH and SP could restore the abnormal expression of α-syn(F), NeuN(G) and VDAC(H). All values are expressed as mean±SEM. All experiments were repeated more than three times individually. Scale bar = 500μm in 100×. \**P* < 0.05, \*\**P* < 0.01, and \*\*\**P* < 0.001.

Consistent with a protective effect of MH on the morphology of neuronal mitochondria, MH treatment also appeared to improve mitochondrial function of the hippocampi in the mice of the ROT+MH group compared to those in the ROT group (Fig. 6B-E). The most commonly used markers for mitochondria functions include malondialdehyde (MDA) and glutathione (GSH), and ATP. As shown in Fig. 6B, ROT increased the MDA level from 1.00 to 1.22 in the hippocampi (*P* =3.0× 10^-3^) of mice, but such a trend was efficiently reversed by MH treatment, as the MDA level of the ROT+MH group decreased to 0.91 (25.41%) (*P* =6.1× 10^-3^) (Fig. 6B). Meanwhile, ROT administration significantly decreased the GSH level in the hippocampus (*P* =2.3× 10^-3^) compared to NC, whereas in the MH+ROT group, the GSH level (*P* =3.8× 10^-2^) was elevated to 152.27, rather higher than the 100.77 in the ROT groups (Fig. 6C). Moreover, in the ROT-induced mice, the ATP level was found to have decreased from 124.02 to 75.61 in the hippocampi (*P* =5.8× 10^-3^) compared to the NC group. In the ROT+MH group, we observed significant elevation of the ATP level from 75.61 to 103.26 (*P* =2.2× 10^-2^) compared to the ROT group (Fig. 6D).

To better characterize the protective effect of MH in the brain, we examined the impact of MH on the homeostasis of mitochondrial marker proteins (Fig. 6E). As shown in Fig. 6F, the level of α-synuclein exhibited by mice in the ROT group increased from 1.06 to 1.35 (*P* =3.5× 10^-3^) as compared to the NC group, whilst expression of α-synuclein decreased from 1.35 to 1.00 in the ROT+MH group (*P* =4.0× 10^-4^) compared to the ROT group. The NeuN level of mice in the ROT group decreased from 0.99 to 0.50 (*P* =2.5× 10^-2^) as compared to the NC group, whereas the ROT+MH groups displayed significant increases (from 0.50 to 0.82) in NeuN expression (*P* =2.5× 10^-2^) as compared to the ROT group (Fig. 6G). The VDAC1 level in the ROT group decreased from 1.49 to 1.06 (*P* =1.4× 10^-2^) as compared to the NC group, whereas the ROT+MH groups displayed significant increases (from 1.06 to 1.26) in VDAC expression (*P* =4.1× 10^-2^) as compared to the ROT group (Fig. 6H). We also examined the impact of SP on homeostasis of mitochondrial marker proteins. The quantitative results are presented in the Supplementary Material (Fig. S8).

To improve our understanding of the mechanism(s) whereby MH improves mitochondrial function in murine hippocampi and neurons, we used qRT-PCR to test a set of genes (*Ndufa12, Cox6b1, Atp5k, Src, Ndufb10* and *Dlgap3*) in the Uni-MH-ROT set that are known to be involved in the pathways pertaining to mitochondrial function and postsynaptic density (Fig. S6A). As shown in Fig. 7A-D, the levels of *Ndufa12*, *Cox6b1*, *Atp5k* and *Src* expression were restored to normal after treatment with MH compared to the ROT group (more detailed results are given in the Supplementary Material, Fig. S6 and Fig. S8).

To better understand how MH might impact synaptic function in the PD mouse model, qRT-PCR was performed to assess the drug‘s effect on the expression of several genes (*Chrna4, Syt11, Cdh8* and *Sncg*) that are known to play a role in synaptic pathways[56–60] in the NC, ROT, MH or MH+ROT groups. As shown in Fig. 7E-H, the expression levels of *Chrna4*, *Syt11* and *Cdh8* were significantly reversed in the ROT+MH group as compared to the ROT group (more detailed results are given in Supplementary Material, Fig. S6 and Fig. S8).

**Fig. 7.**
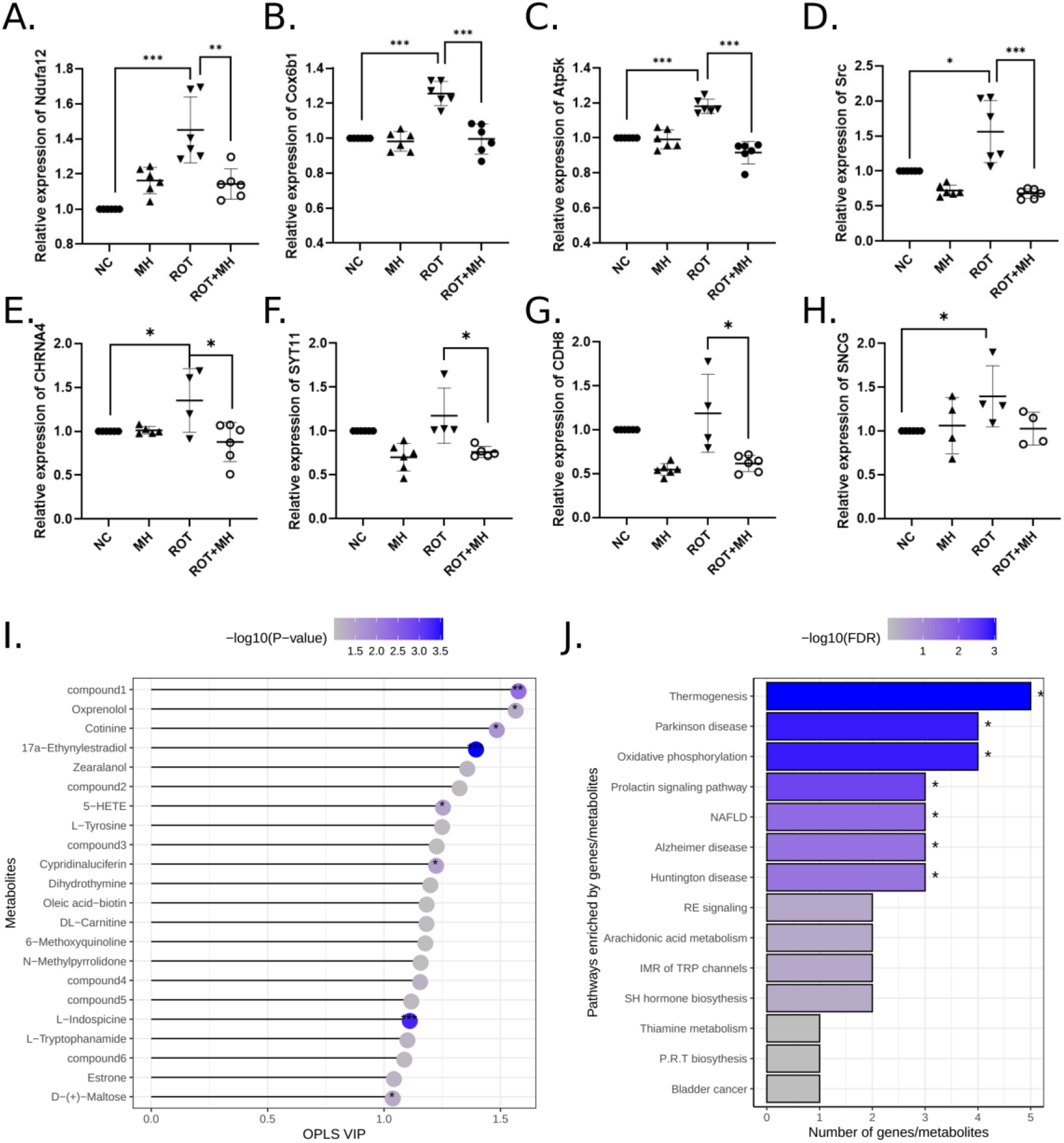
MH regulating genes involved in mitochondrial function, postsynaptic density, and mitochondrial metabolites in a PD mouse model. MH can prevent the dysregulation of lipid oxidation-related genes in brains of mice treated by ROT. Effects of MH on regulating mitochondria-related genes were detected by qRT-PCR in the hippocampus of mice, including (A) *Ndufa12*, (B) *Cox6b1*, (C) *Atp5k* and (D) *Src*. The expression values are given as mean±SEM. The experimental data were taken from more than three independent experiments. MH can prevent the dysregulation of synapse-related genes in brains of mice treated by ROT. Effects of MH on regulating synapse-related genes were detected by qRT-PCR in mouse hippocampus including (E) *Chrna4*, (F) *Syt11*, (G) *Cdh8* and (H) *Sncg*. Expression values are expressed as mean±SEM. The experimental data were taken from more than three independent experiments. \**P* < 0.05, \*\**P* < 0.01, and \*\*\**P* < 0.001. The effect of MH in mitochondrial metabolites of ROT-induced PD primary neuron model. (I) OPLS VIP (variable influence on projection for the orthogonal projections to latent structures model) and Student‘s t-test *P*-values of 22 metabolites whose VIP was higher than 1 and t-test p-values lower than 0.1. \**P* < 0.05, \*\**P* < 0.01; \*\*\**P* < 0.001. (J) KEGG pathway enrichment analysis on 22 metabolites (OPLS VIP>1 and t-test *P* < 0.1) combined with 10 genes (*Chrna4, Syt11, Cdh8, Sncg, Ndufa12, Cox6b1, Atp5k, Src, Ndufb10* and *Dlgap3*) that are indicated in this study as expressed significantly differently in hippocampal regions of the ROT and MH+ROT groups., Compound1: 2-{(3S)-1-[4-(trifluoromethyl)benzyl]-3-pyrrolidinyl}-1,3-benzoxazole Compound2: 2-heptyl-4-hydroxyquinoline-N-oxide Compound3: 5,5-dimethyl-2-{[(2-phenylacetyl)amino]methyl}-1,3-thiazolane-4-carboxylic acid Compound4: *trans*-cinnamoyl beta-D-glucoside Compound5: 6,9-dioxo-11R,15S-dihydroxy-13E-prostenoic acid Compound6: 17beta-hydroxy-5beta-androstan-3-one NAFLD: Non-alcoholic fatty liver disease RE signaling: Retrograde endocannabinoid signaling IMR of TRP channels: Inflammatory mediator regulation of TRP channels SH: Steroid hormone P.R.T biosynthesis: Phenylalanine, tyrosine and tryptophan biosynthesis.

### Identification of a metabolic signature in mitochondria linked to aminoacyl-tRNA and phenylalanine, tyrosine and tryptophan biosynthesis in a MH-treated PD neuronal cell model

To study the metabolic alterations in mitochondria after MH treatment of ROT-induced PD neuronal cells, a global metabolomic analysis was performed in the mitochondria of ROT-induced PD neurons as well as in ROT+MH-treated neuronal cells. Interestingly, eight metabolites measured in whole neuronal cells were significantly (*P* < 0.05) altered in both ROT-induced PD neuronal cells and ROT+MH-treated neuronal cells and with variable influence on projection (VIP) more than 1.0 calculated by the Orthogonal Projections to Latent Structures (OPLS) method (Fig. 7I, Table S12). Among them, the compound of 2-{(3S)-1-[4-(Trifluoromethyl)benzyl]-3-pyrrolidinyl}-1,3-benzoxazole was newly collected in the PubChem database in May 2021, whilst the other seven metabolites have been reported as being related to PD therapeutic or neuroprotective effects[61–73].

We performed KEGG analysis using MetaboAnalytic[74, 75] by combining the 22 metabolites (OPLS VIP >1 and *P* < 0.1) and 10 genes (*Chrna4, Syt11, Cdh8, Sncg, Ndufa12, Cox6b1, Atp5k, Src, Ndufb10* and *Dlgap3*) involved in mitochondrial function or synaptic function, which were expressed significantly differently between the ROT and ROT+MH groups in murine hippocampus. As shown in Fig. 7J, these metabolites and genes are significantly (*P* < 0.05) enriched in seven typical KEGG pathways including mitochondrial function pathways (thermogenesis and oxidative phosphorylation), brain disease pathways (PD, Alzheimer‘s disease and Huntington disease), pathways closely related to PD mechanisms such as NAFLD[76], and the prolactin signaling pathway[77]. Therefore, mechanistically, MH may protect neurons from ROT-induced damage by modulating the function of neuronal mitochondria with an impact on mitochondria-linked metabolic pathways.

## Discussion

We established a method, integrating gene co-expression modules in normal human brain with disease-associated genes or SNPs to identify disease-associated gene co-expression modules that were further used for drug repurposing. iGOLD is able to interpret the impact of individual genes on disease, especially those genes expressed differentially in patients and controls (DEGs) obtained using insufficient samples, which are unable to provide information on the signaling circuitry of disease-associated pathways[78–80]. It can also explain the role of SNPs in regulating gene co-expression modules in normal human brain. These SNPs were discovered as part of large population-based GWAS studies, and their functional roles in regulating gene co-expression modules from normal human brain have remained unclear[81, 82]. When these genes are used for drug discovery, iGOLD determines the drug efficiency not in terms of its target proteins but rather in terms of its ability to restore the normal gene expression profile. This makes iGOLD a powerful tool in drug discovery by dint of its considering multiple genes in one network. This approach can be generally applied to repurposing drugs for other brain disorders simply by connecting any disease-associated genes or SNPs with the gene co-expression modules associated with normal human brain samples.

Our current study suggested MH as a drug candidate for PD. MH is known as a psychostimulant in the nootropic agent group, and is an accepted treatment for traumatic cataphora, alcohol poisoning, anoxia neonatorum, and children’s enuresis[83]. Oral administration of MH to rats in chronic hypoperfusion improved behavioral dysfunction, suggesting an ability of MH to attenuate neuronal damage after ischemia[84]. Another study has investigated the roles of MH in Parkinson‘s disease using a yeast model, and indicated that MH is able to rescue worms expressing α-syn in dopaminergic neurons[85], however, these findings need to be verified in animal model. A previous study has shown that MH may improve muscle tone and brain lipid peroxidation in a rat model[86]. However, no study has been able to show that MH prevents PD by reversing abnormally expressed genes and metabolic factors in certain brain regions. Here, we constructed a rotenone-induced mouse model to validate the biological effects of MH. Many studies have employed rotenone to generate an experimental animal model of PD to mimic the PD-like symptoms, such as motor deficit, cognitive decline and depression[31, 32, 87–89]. We found that MH can ameliorate the PD-related behaviors of mice as measured by the gait change, the sucrose preference test, the forced swim test and the tail suspension experiment. We also provided extensive evidence to show that MH is capable of preventing neuronal death, synaptic damage and the destruction of mitochondria, reducing lipid peroxidation, protecting dopamine synthesis, and reversing the abnormal metabolism of mitochondria, thereby demonstrating that MH operates by improving both mitochondrial metabolism and brain function to ameliorate the most overt symptoms of PD.

Additionally, we found that MH may prevent the further deterioration of Parkinsonian symptoms by improving mitochondrial function, such as impacting the expression of markers for lipid peroxidation and mitochondrial proteins. Moreover, it is widely believed that mitochondrial-associated neurodegenerative diseases involve the perturbation of calcium flux or energy generation[90, 91]. Thus, we measured the ATP levels in the hippocampi of the mice in each group, and noted that MH significantly restores the ATP level in the ROT+MH group to a level comparable to that of the NC group (Fig. 6D). The improved mitochondrial function consequent to MH treatment might also be due, at least in part, to the restoration of normal expression of *Ndufa12*, *Cox6b1*, *Atp5k* and *Src* genes in the ROT-induced PD mouse model. *NDUFA12* gene has been shown to encode a key member of the mitochondrial respiratory chain[92, 93]. Low expression of the *Cox6b1* gene has been associated with Alzheimer’s disease[94]. In similar vein, *ATP5K* is known to be involved in mitochondrial ATP synthesis-coupled proton transport[95].

We also investigated the impact of MH on mitochondria by means of mitochondrial metabolomics, and disclosed several specific metabolites that were regulated by MH, suggesting that MH may influence mitochondrial function by reprogramming metabolic pathways. Understanding drug-metabolite associations is crucial for research into pharmacoepidemiology and for improving drug efficiency[96]. One recent study has demonstrated that metabolic abnormalities can alter neuronal excitability in the brain[97]. We found that MH treatment can restore normal levels of several metabolites associated with PD pathogenesis including 5-HETE and L-indospicine. For example, 5-HETE (OPLS VIP = 1.250 and t-test *P* = 3.9 × 10^-2^) has been reported as a biomarker of oxidative damage in PD^61^; 5-HETE interacting with *SRC* regulates the *TRPV1* gene, which has been reported to be associated with PD development[98–103]. Another compound, L-indospicine (OPLS VIP=1.107, *P* = 5.0 × 10^-4^), has been reported to be a potent inhibitor of arginase that can cause a shift in L-arginine metabolism to the NOS pathway^64^ closely related to PD development[104–106].

Damage to synaptic plasticity is also known to be related to the onset and progression of both the motor and cognitive symptoms of PD[107]. Previous studies have employed immunohistochemistry to investigate the protective potential of MH in relation to synapses[108, 109]. However, these studies could not determine the true length and numbers of synapses. In order to confirm the protective action of MH on synapses, we performed *in vitro* and *in vivo* experiments as well as cluster analysis to demonstrate that MH can protect synapses in terms of synaptic length. Additional q-PCR experiments indicated that MH treatment of ROT-induced PD primary neurons restores normal expression of the *Chrna4*, *Syt11* and *Cdh8* genes. These genes have been previously shown to encode proteins with functions pertaining to synaptic function[57-59, 110-113]. Thus, our study supports the view that MH may protect synapses by impacting the pathways in which both mitochondria-related genes, and metabolic factors such as maltose and cotinine, are involved.

Our studies do however have several limitations. Firstly, the tissue chip can only interrogate part of the synapses, and is unable to fully observe the protective effect of MH on the murine synapse. New technology for observing whole synapses will be required to confirm the protective effect of MH. Secondly, the CMAP database only includes a limited number of drugs, which may hinder the identification of more effective drugs for repurposing. Thirdly, the effectiveness of the drugs themselves still requires further supporting evidence from clinical studies.

In conclusion, this study revealed MH as a potential drug candidate for PD. Subsequent experiments indicated that MH is able to improve PD-related behavior and protect neurons by regulating mitochondrial-related genes, synaptic pathways and metabolite pathways. Thus, it would appear that MH may help to arrest the progressive deterioration of Parkinsonian symptoms.

## Acknowledgments and Funding

The work was funded by the National Key Research and Development Program of China (2020YFB0204803), the Natural Science Foundation of China (81801132 and 81971190), Guangdong Key Field Research and Development Plan (2019B020228001 and 2018B010109006), and Natural Science Foundation of Guangzhou (2021A1515010256). Medical and Health Science and Technology Plan of Longgang Shenzhen (No. LGKCYLWS2022019).

## Materials and Methods

### Study design

Here, we designed a computational architecture, iGOLD, for drug repurposing. This approach involved the construction of gene co-expression modules of normal human brain by applying weighted gene co-expression network analysis (WGCNA)[114] and DiffCoEx[115] and analyzing gene expression data of 1,231 brain samples from ten brain regions of healthy human. Then, iGOLD was used to identify the modules enriched in PD-associated genes and PD-associated SNPs by employing 11 datasets encompassing PD-associated genes, SNPs and DEGs between PD and controls. The identified modules were evaluated in relation to their expression conservation in brain samples across ethnicities, brain regions, and disease stages of PD by ModulePreservation, a function in the WGCNA R package[116]. This analysis was based upon seven datasets with sample sizes ranging from 4 to 57. The highly conserved modules were used for drug repurposing by CMAP[38, 39]. The drug candidates were ranked by their connectivity scores. From them, we selected those ranked in the top 15 (Table S9) and having the ability to pass through blood brain barrier for further validation. The source code of iGOLD and related data used in this study are available at https://github.com/fanc232CO/iGOLD_pipline.

The experimental validations of the drug effects were conducted in primary neurons and a mouse model. The primary neurons were obtained from the hippocampi of mice on postnatal days 0-3. We used Rotenone (ROT) to treat the primary neurons as described previously[117] since ROT has been shown to induce PD-like symptoms in human[27]and animal models[28, 29]. The effectiveness of drugs in protecting neuronal damage was evaluated by immunofluorescence marking the tubulin of synapses and primary neurons, and a mitochondrial fluorescent probe for mitochondrial morphology. The protective effects of the drug on mitochondrial functions were tested by determining the malondialdehyde (MDA) level, the reduction in the glutathione (GSH) level, ATP levels and mitochondrial proteins. The regulatory effects of drugs on mitochondrial metabolites were assessed in murine primary neurons.

The mouse model was constructed using ROT-induced C57B/L male mice (*n* = 86, 8 weeks old) (Supplementary Material). We used PET/CT imaging to examine glucose metabolism in the brains of mice (Supplementary Material). The protective ability of the drugs on cranial nerve damage was evaluated by immunohistochemistry of NeuN (in the DG, DG2 and CA1), NeuN-negative cells in the DG structure of the hippocampus, NeuN-negative cells in the DG2 structure of the hippocampus, NeuN-negative cells in the CA1 structure of the hippocampus, and TH in the hippocampus. Expression of mitochondrial-related proteins was measured by western blotting for α-syn(F) and NeuN(G) in the murine hippocampi. RNA-seq and qRT-PCR were used to measure the expression of mitochondrial-related genes in the hippocampi of mice. The effectiveness of the drugs was further examined in terms of their influence on the PD behaviors of mice including the footprint test, sucrose preference test, and forced swim test.

### Gene expression data used for identifying modules associated with PD

Gene expression data were obtained from the publicly available GEO dataset[118]. From GEO, we downloaded eight gene expression datasets [one for co-expression module building, five for module conservation analysis, and two for enrichment analysis of Parkinson’s disease (PD)-associated DEGs].

The co-expression modules were constructed by WGCNA analysis on the GSE60862[35–37], a gene expression dataset obtained from the platform of Affymetrix Human Exon 1.0 ST Array. The samples covered ten brain regions, including cerebellar cortex, frontal cortex, occipital cortex, temporal cortex, hippocampus, putamen, thalamus, medulla, white matter and substantia nigra, from 1,231 individuals of European descent collected by the UK Brain Expression Consortium (UKBEC).

The module conservation analysis was performed on six GEO datasets, GSE131617[41, 119], GSE23290[42], GSE34516[43], GSE51922[44], GSE18838[45] and GSE34865 (https://www.ncbi.nlm.nih.gov/geo/query/acc.cgi) which were obtained from the platform of the Affymetrix Human Exon 1.0 ST Array. These datasets covered multiple ethnicities, brain regions, tissues, Braak stages, and PD disease status. In detail, GSE131617 includes transcriptome data from 213 post-mortem brain tissue specimens (= 71 subjects × 3 BRs), which covered three brain regions (entorhinal, temporal and frontal cortices) of 71 Japanese brain-donor subjects in four Braak stages[120–122] (0, I–II, III–IV, V–VI). GSE23290 included putamen tissues from the 8 idiopathic PD (IPD) patients, 3 LRRK2-associated PD (G2019S mutation) patients, 5 neurologically healthy controls, and one asymptomatic *LRRK2* mutation carrier[42]. GSE34516 included the locus coeruleus post-mortem tissues from idiopathic PD (IPD) and LRRK2-associated 6 European PD patients[43]. GSE51922 is built on the RNA profile of IPSC-derived dopaminergic neurons from idiopathic and genetic forms (LRRK2) of PD^43^. GSE18838 included peripheral blood collected from 18 PD patients and 12 healthy controls[45]. GSE34865 included gene expression data of substantia nigra samples from 57 healthy adults. Details of these datasets can be found in Table S7.

GSE8397[40, 123] were downloaded for generation of PD-associated differentially expressed genes (DEGs). GSE8397 is built on the gene expression of substantia nigra split into medial and lateral portions, and frontal cortex from 24 PD patients and 15 controls. Gene expression was accessed through two platforms, GPL96 and GPL97[40, 123]. Details of these datasets can be found in Table S6.

### Conservation analysis of co-expression gene module

Conservation of modules was estimated by modulePreservation through calculation of *Z_summary_*. The *Z_summary_* is determined by estimating the density and connectivity of the test modules and reference module. Briefly, the calculation of *Z_summary_* is based on permutation tests to assess the mean and variance of Z statistics under the null hypothesis of no relationship between the module assignment in reference and test modules. The reference gene expression data in this study were derived from seven GEO databases (Table S7), while the test modules are the PD-associated modules suggested by iGOLD. Modules with *Z_summary_*scores above 10 were interpreted as being highly conserved, *Z_summary_* scores between 2 and 10 were deemed to be moderately conserved, whilst *Z_summary_* scores below 2 were regarded as incompletely conserved.

### Stratified LD score regression (sLDSC) analyzing PD-associated SNPs

sLDSC analysis[124] was conducted using the parameters and pipelines provided by tutorials in LDSC (https://github.com/bulik/ldsc/wiki). First, we mapped all SNPs to the co-expressed modules if they were within 10kb of the locations of exon probes. The LD score and heritability were calculated for each co-expressed module. The enrichment of the SNPs in the co-expression modules was defined as the summation of SNP heritability divided by the number of SNPs in that module. Standard errors of the SNP enrichment in the co-expression modules were estimated by a block jackknife[125] and were further used to evaluate Z-scores, *P*-values and false discovery rates (FDRs) of the SNP enrichment in the co-expression modules[73].

### Construction of co-expression networks for normal human brain

The GSE60862[35–37] dataset was used to construct the gene co-expression networks. The genes expressed in ten brain regions were grouped into co-expression modules by two different approaches, consensus weighted gene co-expression network analysis (WGCNA)[114] and DiffCoEx[115]. The WGCNA was applied to detect co-expression modules common to all ten brain regions (consensus co-expressed modules, CCM). DiffCoEx was used to identify gene modules specifically expressed in each of the ten brain regions compared to the other nine brain regions (specific co-expressed modules, SCM).

The function, blockwiseModules in WGCNA, was utilized to construct the co-expression modules as previously described[126, 127]. The parameters were set as follows, β = 7 (chosen based on the scale free topology criterion r^2^ > 0.8), minModuleSize = 30, mergeCutHeight = 0.25, maxBlockSize = 6000, and corType = ‗pearson‘. For each pair of genes, the topological overlap matrix (TOM) was calculated and scaled based on the adjacency matrix. The component-wise minimum of the TOMs in each brain region was then extracted to generate a consensus TOM. This TOM was clustered by using the average hierarchical clustering method to obtain a consensus TOM defining it as 1-consensus TOM according to the difference of genetic connectivity. A consensus co-expression module was defined as a branch of a cluster tree generated by a dynamic tree cut.

The differential co-expression network analysis was carried out by the DiffCoEx method in R software as previously described[115]. To identify gene co-expression differences between transcripts from the substantia nigra brain region and transcripts from the other nine brain regions, we used the function of DiffCoex based on the WGCNA framework by calculating a TOM generated from a matrix of adjacency differences between these brain regions.

### Module conservation analysis

The conservation of the association between the gene co-expression modules and PD was evaluated according to the enrichment of the modules in DEGs of PD patients and controls from five different sources[41–45, 119]. These datasets were all obtained from the GPL5175 platforms, and the samples in the datasets were divided into healthy control and PD groups. The samples in this dataset are accompanied by information on Braak stages indicating the disease severity[120–122]. These samples were partitioned in terms of two ethnicities, three brain tissues, five brain regions and six PD disease states. Further information is provided in Table S7. The degree of module conservation was estimated by modulePreservation, a function in the WGCNA R package[116].

### Enrichment analysis of PD-associated genes

We validated the association between PD and CCM and SCM modules by the enrichment analysis of PD-associated genes and PD-associated SNPs. The PD-associated genes were obtained from DisGeNet[128–130] (https://www.disgenet.org/home/). DisGeNET covers the full spectrum of human genetic diseases as well as normal and abnormal traits. The currently released version of DisGeNET includes more than 24,000 different genetic diseases and traits, 17,000 genes and 117,000 genomic variants[130]. As shown in Table S3, searching under the term ―Parkinson disease‖ (UMLS CUI: C0030567) allowed the collation of six types of PD-associated genes by the DisGeNet database, of which ―CausalMutation‖ was filtered out before enrichment analysis because it contained only one gene. Gene enrichment analysis in the remaining five gene sets was evaluated by means of the single-tailed Fisher‘s exact test, and further adjusted by the false discovery rate (FDR).

### Enrichment analysis of genes expressed significantly differently in PD patients and controls

Differentially expressed genes (DEGs) between PD patients and controls were obtained by analyzing gene expression data from GEO with access ID GSE8397[40, 123] (Table S6). In GSE8397, samples from the whole substantia nigra (combination of samples from lateral and medial substantia nigra regions) were used for DEG analysis. The DEG analysis was performed by means of the GEO2R tool[131, 132] in the GEO website (https://www.ncbi.nlm.nih.gov/geo/). DEGs were selected with the fold change (FC) threshold of 1.2 and adjusted *P*-value threshold of 0.05. Enrichment was evaluated by single-tailed Fisher‘s exact test adjusted by the False Discovery Rate (FDR).

### Drug discovery by CMAP

First, the DEGs were generated by analyzing gene expression data (GSE8397) of substantia nigra samples obtained from the platform of GPL96 (Affymetrix) (15 healthy controls and 24 PD). The overlapping genes between these DEGs and the genes in PD-related modules were extracted and mapped to the GPL96 probe, and then delivered to the Connectivity Map (CMAP)[38, 39] (https://portals.broadinstitute.org/cmap/) to predict potential PD drug candidates. We ranked the output drug candidates by their connectivity scores. When the connectivity scores were close to-1, the drugs were deemed to have strong potential to restore the normal gene expression profile of the PD-associated genes. In this study, drug candidates were selected for further analysis if their connectivity scores were lower than-0.8.

### Neuronal cell culture

The hippocampal primary neurons were obtained from mice on postnatal days 0-3. Hippocampal neurons were plated on poly-D-lysine (Sigma)-coated chamber slides and Six-hole plates for 2 hrs to allow neurons to adhere. The cultured neurons were maintained with complete culture medium composed of B27 supplement (Gibco, USA), L-glutamine (Life Technologies) and Neurobasal-A medium (DMEM/F12) (Gibco, USA) at 37 ℃ in a 7% CO_2_ incubator for 7 days to ensure the growth of nerve synapses.

### Construction of neuron models

The primary neurons were randomly divided into 6 groups: the normal control (NC) group, the ROT-induced group, the SP-treated group, the MH-treated group, the ROT+SP treated group and the ROT+MH treated group.

To construct the ROT-induced group, ROT with concentration 400 nM was applied directly to the culture medium for 24 hrs. To create ROT+SP(2μM) and ROT+MH(10 μM) groups, we pretreated murine primary neurons with SP or MH for 2 hrs, and then used 400 nM rotenone (ROT) to treat the neuronal cells for 24 hrs. The SP and MH groups of the murine primary neurons were treated by SP or MH for 2 hrs. We then used immunofluorescence marking the tubulin of synapses and primary neurons to evaluate the cell damage from ROT.

### Construction of mouse models

C57B/L male mice (*n* = 86, 8 weeks old) were randomly assigned into six groups, namely the control group (NC), the Sodium phenylbutyrate (SP) group, the Rotenone (ROT) group, the ROT+SP group, the Meclofenoxate-hydrochloride (MH) group, and the ROT+MH group. The mice in the NC group received dimethylsulfoxide (DMSO) (olive oil only); the mice in the SP group were treated with SP (300 mg/kg bw/d; intraperitoneal [i.p.]) for 4 consecutive weeks; the mice in the ROT group were given rotenone (1 μg/g bw/d; i.p.) for 3 consecutive weeks; the mice in the ROT+SP group received SP prophylaxis (300 μg/g bw/d; i.p.) for 1 week followed by ROT (1 μg/g bw/d, i.p.) from 1 week onwards for the next 3 consecutive weeks; the mice in the MH group were administered with MH (50 μg/g bw/d; i.p.) for 4 consecutive weeks; the mice in the ROT+MH group received MH prophylaxis (50 μg/g bw/d; i.p.) for 1 week followed by ROT challenge (1 μg/g bw/d, i.p.) from 1 week onwards for the next 3 weeks.

### Body weight and food intake of ROT-induced PD mouse model

In this study, 36 C57BL/6 mice were randomly divided into 6 groups that were treated with DMSO (NC), ROT, SP, MH, ROT+SP and ROT+MH, each group containing six mice. Administration of SP and MH did not elicit any behavioral alterations during the experimental period, nor were any significant changes in food intake evident. Mice treated with ROT showed no decrease in body weight during the treatment period. Mice in other groups did not exhibit any significant decrease in body weight during the treatment period nor any significant changes in food intake, except for the ROT group (Table S8).

### Hippocampus sample collection

From each group of mice, more than three hippocampi were collected. The mice were sacrificed by anesthetization, and their brains were extracted within 24 hrs after the last injection of ROT. The hippocampi were then isolated under a microscope.

### Biospecimen collection

The striatum brain regions of mice were separated and processed to obtain both cytosolic and mitochondrial fractions, after they were sacrificed by anesthetization, within 2 hrs. The biochemical investigations were conducted and 4–5 murine striatums (from each group) processed for histopathological examination.

### RT-PCR

Total cell RNA extraction was performed using TRIzol (Invitrogen) and reverse transcribed according to the manufacturer‘s protocol (Takara). qRT-PCR was carried out using SYBR Premi qRT-PCR ExTaq™ II (Takara) and analyzed on a Bio-Rad CFX96 real time PCR cycler (BioRad, Netherlands). The primer sequences are given in Table S13. Differences in mRNA expression were calculated by means of the formula N = (2)^-ΔΔCT[133].

### Immunofluorescence

Samples were washed three times with 0.01 M PBS and fixed with 3.7% paraformaldehyde for 15 min at room temperature. The samples were then permeabilized in 0.5% Triton X-100 for 3 min and blocked with 3% goat serum albumin for 30 min prior to incubation with a primary antibody, namely, anti-α-synuclein, mouse anti-TH and anti-NeuN, at dilutions of 1:100 overnight at 4℃. Secondary antibodies, anti-rabbit or anti-mouse (Tianjin Sungene Biotech Co., China) at 1:200 dilution, for 1 hr at room temperature. Nuclei were stained with DAPI.

### Immunohistochemistry

The slides were washed twice for 15 min in 0.01 M PBS, and proteinase K was added to the tissue and incubated at 37°C for 5 min. This step was followed by quenching for 10 min in a solution of methanol containing 30% hydrogen peroxidase and further incubating for 1 h in blocking solution containing 5% normal goat serum and 1% Triton X-100 in 0.01 M PBS. After blocking, the slides were incubated overnight in rabbit anti-caspase-3 antibody (Catalog Number:10842-1-AP, Proteintech), anti-α-synuclein antibody (Catalog Number:10842-1-AP, Proteintech, China), anti-TH antibody (Catalog Number:25859-1-AP, Proteintech, China) and anti-NeuN antibody (Catalog No.: A19086, ABclonal, China) diluted 1:100 in blocking solution. Following incubation with primary antibody, the sections were incubated for 2 h in biotinylated goat antirabbit secondary antibody diluted 1:500 in 0.01 M PBS and subsequently incubated with ABC reagents (Standard Vectastain ABC Elite Kit; Vector Laboratories, Burlingame, CA, USA) for 20 min in the dark at room temperature. The sections were washed twice with 0.01 M PBS and incubated in 3,3′-diaminobenzidine tetrahydrochloride (DAB); sections were washed with distilled water, dehydrated in graded ethanol (70%, 85%, 95% and 100%), placed in xylene and cover slipped using mounting medium. We then analyzed and counted the active caspase-3 positive cells in the dentate gyrus (DG), dentate gyrus2 (DG2) and cornu ammonis (CA1) regions of the hippocampus using ImageJ program analysis.

### Mitochondrial Fluorescent Probe Staining Analysis

Mitochondrial staining was performed with the mitochondrial probe MitoTracker Red CM-H2XRos (Invitrogen, USA) according to protocols provided by the manufacturer. After being washed with 0.01 M PBS, the cells were counterstained with DAPI for 10 min and imaged with an Olympus BX63 microscope (Olympus, Japan).

Neurons from differentiated groups were stained with MitoTracker Deep Red (200 ng/ml) (Yeasen, Shanghai, China) for mitochondria for 60 min, then fixed with 4% paraformaldehyde for 15 min and permeabilised by Triton X-100 at 0.04% as previously described[134]. All cells (nuclei) were stained with DAPI (4′,6-diamidino-2-phenylindole, 1 µg/ml). Images were obtained using an Olympus BX63 microscope (Olympus, Japan). Quantification and analysis of neuronal network was performed using Image J software.

### Footprint test

Gait patterns of mice were assessed by the footprint test as described previously[135]. The apparatus comprising an open field (60 × 60 × 40-cm), in which a runway (4.5 × 40 × 12 cm) was arranged to lead out into a dark wooden box. Gait parameters were measured by wetting fore paws and hind paws with commercially available non-toxic colored inks and allowing the mice to trot onto a strip of paper on the runway. Pawprints made at the beginning and the end of the run were excluded. Various gait parameters such as stride length (Differences in the forward distance between each fore paw and hind paw footprint with each step), stride width (lateral distance between opposite left and right fore paw and opposite left and right hind paw) and foot direction, were measured.

### Sucrose Preference Test

The mice were housed individually and were first trained to adapt to sugary drinking water in a quiet room by putting two water bottles in each cage. Both bottles were filled with 1% sucrose water. The mice were tested with respect to sucrose preference using the following process: (1) the mice were prevented from drinking for 24 hrs before administration of the Sucrose Preference Test; (2) each mouse was given a pre-quantified bottle of 1% sucrose water and a bottle of distilled water; (3) the position of the two bottles of water was changed every 12 hrs; (4) the two bottles of water were taken and weighed after 24 hrs to calculate the consumption of sucrose water, distilled water and total liquid consumption for each mouse. Sucrose water preference (%) = (sucrose water consumption/total liquid consumption) × 100%.

### Forced Swim Test

The mice were placed individually into an open cylindrical container (10 cm diameter, 30 cm height), containing water (25 ± 2°C) to a depth of 20 cm. Each mouse was forced to swim for 6 min, and the total duration of immobility in seconds was measured during the last 4 min. The water was changed after each animal experiment was finished. After the experiment, the mice were wiped with a towel until their fur was dry. The immobility time was defined in terms of the absence of escape-oriented behavior.

### PET/CT imaging and data analysis

The PET/CT imaging was performed on 36 C57B/L mice that were divided into NC, SP, ROT, ROT+SP, MH and ROT+MH groups. Each group contains 6 mice. All PET imaging studies were performed on a Biograph TrueV (Siemens Healthcare) scanner. This PET/CT device is equipped with a 64-slice spiral CT component. After a 3-h fasting period, mice were injected in the tail vein with a solution of ^18^F-FDG (16–32 MBq) or ^18^F-FLT (32–37 MBq). The volume of the syringes was always kept below 0.2 mL in order to meet the requirements of our ethics committee. To minimize muscle and brown fat uptake in the case of ^18^F-FDG imaging, animals were kept anesthetized under warming lights for a 20-min period after injection. Animals were imaged simultaneously in groups of three with the PET/CT scanner; one was placed at the center of the field of view (FOV) whilst the two others were placed on each side of the central animal, at a 5-cm and at a −5-cm radial offset.

The PET/CT was initiated with a CT scan acquired with the following parameters: 80 mA, 130 kV, pitch 0.8, and 64×0.6mm collimation. Then, an emission scan was obtained in 3D mode. PET images were reconstructed in a reduced FOV (35 cm) applying a scaling factor of 2. Images were reconstructed with an algorithm which models point spread function of the scanner and leads to a 2.2 mm spatial resolution at the center of the FOV. The following parameters were used: six iterations, 16 subsets, no filtering, and a matrix size of 3362, resulting in a 1.02×1.02×1mm voxel size. Scatter and attenuation corrections were applied.

Brain activity was obtained from a volume of interest (VOI) encompassing the entire brain. The VOI was determined by means of an isocontour, which was set so that the VOI matched the apparent brain volume on PET and CT images. When discordance was encountered between the PET metabolic volume and the CT volume, the VOI was drawn according to CT images, so that PET/CT images could be compared to *ex vivo* counting of the entire hippocampus for which the entire brain was harvested, irrespective of the presence of non-viable areas. Data were analyzed using Statistical Parametric Mapping software (SPM 12, The Wellcome Trust Center for Neuroimaging, London, UK).

Relative [18F] FDG uptake images were analyzed by using microQ (Siemens/Concorde Microsystems, Knoxville, TN, USA). Subsequently, we utilized a voxel-by-voxel approach to obtain maximal use of information without *a priori* knowledge, using IRW (Siemens/Concorde Microsystems, Knoxville, TN, USA). In short, we used a flexible factorial design depending on time point (after all treatment) and group (NC, SP, ROT, SP + ROT, MH and MH+ROT), as previously described (1, 2). T-maps were interrogated at a *P_height_* ≤ 0.005 (uncorrected) peak level and extended threshold of kE > 200 voxels (1.6 mm^3^)[136]. Only significant clusters with *P_height_* < 0.05 (corrected for multiple comparisons) were retained.

### ELISA analysis

α-synuclein was quantified in the striatum region of mouse brain utilizing the Enzyme-linked Immunosorbent Assay (ELISA) Kit from Invitrogen (Cat No. KHB0061) following the manufacturer‘s instructions.

### Western blot (WB) analysis

For western blots, cells were extracted in RIPA buffer (Sigma-Aldrich, USA), separated by 10% SDS-PAGE and transferred onto a polyvinylidene difluoride membrane (Millipore, USA). After blocking with 5% skimmed milk, the membrane was incubated with specific primary antibodies directed against α-synuclein, TH and GAPDH (Abcam plc, USA]) at a dilution of 1:1,000 overnight at 4℃. The protein expression levels were normalized with GAPDH. The membrane was incubated with horseradish peroxidase-conjugated anti-rabbit secondary antibody for 1 hr at 37℃.

### Measurement of oxidative stress markers

The markers of oxidative impairment were studied in the cytosol. MDA (malondialdehyde) contents were quantitatively detected using the Lipid Peroxidation MDA-Assay Kit (Beyotime). Unit weight of MDA was calculated by a MDA standard curve measured at 532 *nm*. MDA was considered as a biomarker of lipid peroxidation in tissues and organs[137].

### Adenosine triphosphate (ATP) measurement

ATP was measured using ATP assay kits (Beyotime, Shanghai, China). After being diluted with dilution buffer, ATP detection reagent was added to a 96-well plate. After homogenization followed by centrifugation at 12,000 g at 4 ℃ for 5 min, the samples were added into the wells and mixed with the detection solution. Then, the levels of ATP were measured with a SpectraMax M5 microplate reader (Molecular Devices, San Jose, CA, USA). The ATP content was normalized to the ATP protein content on the basis of the standard curve.

### Quantitative analysis of GSH levels

Estimation of GSH: Glutathione (GSH) was measured in the supernatant of hippocampus tissue in mouse brain. GSH level was measured with the enzyme-linked immunoassay (ELISA) kit from Elabscience (Bethesda, MD) according to the manu-facturer′s instruction.

### Sample preparation and library preparation for transcriptome sequencing

Hippocampal RNA was extracted by TRIzol. RNA quantification and qualification were evaluated as follows: (1) RNA degradation and contamination were monitored on 1% agarose gels; (2) RNA purity was checked using the NanoPhotometer® spectrophotometer (IMPLEN, CA, USA); (3) RNA integrity was assessed using the RNA Nano 6000 Assay Kit of the Bioanalyzer 2100 system (Agilent Technologies, CA, USA).

A total of 1 µg RNA per sample was used as input material for the RNA sample preparations. Sequencing libraries were generated using NEBNext® UltraTM RNA Library Prep Kit for Illumina® (NEB, USA) following the manufacturer‘s recommendations; index codes were added to attribute sequences to each sample. Briefly, mRNA was purified from total RNA using poly-T oligo-attached magnetic beads. Fragmentation was carried out using divalent cations under elevated temperature in NEBNext First Strand Synthesis Reaction Buffer (5×). First strand cDNA was synthesized using a random hexamer primer and M-MuLV Reverse Transcriptase (RNase H-). Second strand cDNA synthesis was subsequently performed using DNA Polymerase I and RNase H. Remaining overhangs were converted into blunt ends via exonuclease/polymerase activities. After adenylation of 3‘ ends of the DNA fragments, NEBNext Adaptor with hairpin loop structure were ligated in preparation for hybridization. In order to select cDNA fragments of preferentially ∼250-300 bp in length, the library fragments were purified with the AMPure XP system (Beckman Coulter, Beverly, USA). Then, 3 µl USER Enzyme (NEB, USA) was used with size-selected, adaptor-ligated cDNA at 37℃ for 15 min followed by 5 min at 95 °C before PCR. Then PCR was performed with Phusion High-Fidelity DNA polymerase, Universal PCR primers and Index (X) Primer. At last, PCR products were purified (AMPure XP system) and library quality was assessed on the Agilent Bioanalyzer 2100 system.

### Clustering and transcriptome sequencing

The clustering of the index-coded samples was performed on a cBot Cluster Generation System using TruSeq PE Cluster Kit v3-cBot-HS (Illumina) according to the manufacturer‘s instructions. After cluster generation, the library preparations were sequenced on an Illumina Novaseq platform and 150 bp paired-end reads were generated.

### Metabolite extraction and UPLC–MS/MS analysis

The mitochondrial metabolite tests were performed on murine primary neurons. Global metabolic profiles were obtained from the cells using the Metabolon Platform. The principle of the Metabolon Platform has been previously described^138-140^. Approximately 5×10 ^6^ primary neuronal cells were involved in the test. These cells were extracted from mice at postnatal days 0-3 as described above. The primary neuronal cells were extracted and cultured for 7 days, treated with MH or DMSO for 2 hrs, and then with ROT or DMSO for 24 hrs, to form a PD cell model, which was used for the experiments described below. To lyse the cells, 1,120 μL lysis system (800μl pre-cooled methanol + 320 μL ice water) was added to the wells of the six-well plate; the cellular metabolites were placed in a 2ml EP tube, and 800μL pre-cooled chloroform was injected into the tube prior to vortexing for 15 min. The mixture was then centrifuged at 12,000 rpm for 15 min before being separated into supernatant and precipitate, which were transferred to other microtubes. The supernatant was then dried by a continuous flow of nitrogen gas to render it solid; the solid was re-dissolved with 100 µL acetonitrile-water (1:1), which was centrifuged at 14,000 rpm for 5 min; a mixture of supernatants of the sample (10 µL) was transferred to a quality control (QC) vial. The samples were kept on ice throughout the procedure unless centrifuged. Ratios above were all according to volume.

For UPLC-MS/MS analysis, each sample was reconstituted by a methanol solution with density 80% (80% methanol: 20% water) (by an 80-μL volume of methanol with 20-μL H_2_O). The methanol solution was centrifuged at 12,000 g for 10 min. The samples were then prepared for liquid chromatography-mass spectrometry. Briefly, the samples were injected into a Waters ACQUITY UPLC BEH Amide Column (2.1×100mm, 1.7µm) at column temperature, 40 ℃, with a flow rate of 0.35 mL/min. The mobile phase including phase A and B (Phase A: 95:5 (acetonitrile: water) containing 10mM ammonium format, 0.1% formic acid; Phase B: 50:50 (acetonitrile: water) containing 10mM ammonium format, 0.1% formic acid). The gradient elution ratio being shown in Supplementary Table S14. Subsequent analyses were performed using ThermoScientific Ultimate 3000 UPLC couple with Orbitrap Exploris 480 MS from Sun Yat-Sen Memorial Hospital. Identification of known chemical entities was based on comparison with metabolomic library entries of purified standards. Each biochemical was rescaled to set the median equal to 1. Values for each sample were normalized by Bradford protein concentration.

### Metabolomic instrumentation and analytical conditions

LC-MS/MS was used for detection, with at least 3 replicates for each experimental condition.

### Binding target prediction for MH and SP

Binding targets of MH and SP were predicted using the tool DStruBTarget[48]. The inputs of DStruBTarget are structures of MH and SP, which were obtained from the PubChem database (https://pubchem. <u>ncbi.nlm.nih.gov/</u>) in the format of 3D sdf (compound CID:4039 for MH and 5258 for SP). The parameters of DStruBTarget were employed as the defaults in the prediction. The top ten binding targets with the highest scores were analyzed in this study. The structures of the protein-drug complexes were predicted by SwissDock (http://www.swissdock.ch/)]. The protein structures were downloaded from the PDB database. The PDB ID of *CNR1* predicted as bound by MH is 5TGZ^142^. The PDB ID of *CASP1* predicted as bound by SP is 1BMQ. The structure of the drug-protein complex interaction was visualized using open-source pymol (version 1.7.x) (http://www.pymol.org).

## Data Availability

The source code of iGOLD and related data from this study are available at https://github.com/fanc232CO/iGOLD_pipline.

## Competing interests

The authors report no competing interests.

## Author contributions

H. Zhao, R. Hu, and YD. Yang designed the study. H. Zhang and C. Fan performed the analyses with assistance from H. Zhao. H. Zhang, L. Li, F. Liu, and L. Ma contributed to experimental design, computational analyses, immunohistochemistry experiments, cell culture study, neuronal nuclei isolations, metabolomics experiments, mitochondrial related experiments and behavioral analyses in mice. H. Zhang, C. Fan, YH. Yang, D.N. Cooper and H. Zhao wrote the manuscript. YD. Yang, YH. Yang, D.N. Cooper, and H. Zhao supervised the study. All authors discussed the results and interpretation, and contributed to the final version of the paper.

## Notes

### Competing Interest Statement

The authors have declared no competing interest.

### Summary of Updates

We had corrected the author information

