## supplementary material for "Repurposing the Memory-promoting Meclofenoxate Hydrochloride as a Treatment for Parkinson‘s Disease through Integrative Multi-omics analysis"

**Results**

**Application of iGOLD in the discovery of PD-associated co-expression modules**

iGOLD identified significant enrichment of PD-associated genes in 12 CCM modules and 34 SCM modules (Chi-square test *P* < 0.05) (Fig. S2), whereas 4 CCM modules and 32 SCM modules were significantly enriched with genes within 10 bp of the PD-associated SNPs (Fig. S2). Further, iGOLD identified 5 CCM modules and 22 SCM modules that were significantly enriched in PD-associated SNPs according to linkage disequilibrium (LD) score regression (LDSC)[1](Fig. S2).

We examined the enrichment of DEGs in five modules, M3, BR7M4, BR9M3, BR6M3 and BR3MR (Fig. S4). The proportions of GPL96- and GPL97-DEGs in BR7M4 were 43% and 33%, respectively, which is significantly (Chi-square test FDR = 6.42$\times$10^-21^, 2.28$\times$10^-20^, 2.28$\times$10^-20^, and 3.83$\times$10^-16^ for GPL96, and Chi-square test FDR = 2.45$\times$10^-24^, 2.54$\times$10^-23^, 1.05$\times$10^-19^ and 8.35$\times$10^-19^ for GPL97, respectively) higher than the DEGs in M3 (21% and 13%), BR9M3 (21% and 12%), BR6M3 (21% and 14%) and BR3M2 (23% and 13%) (Fig. S4). We also compared BR7M4 with the other three modules, BR9M3, BR6M3 and BR3M2, that were specifically expressed in substantia nigra, thalamus, putamen and hippocampus, respectively. As shown in Fig. 2B, BR7M4 was significantly enriched with PD-associated SNPs from two sources, GCST007780 according to sLDSC analysis and GCST010765 according to Chi-square test, and was also enriched in PD-associated genes collected by DisGeNet database from five different sources (AlteredExpression, Biomarker, CausalMutation, GeneticVariation, Posttranslational Modification and Therapeutic). By comparison, BR6M3, BR9M3 and BR3M2 did not show any significant enrichment in PD-associated genes collected by DisGeNet database from all the five sources. Additionally, BR3M2 was not enriched with SNPs from GCST007780.

**iGOLD can identify candidate drugs for PD by targeting the BR7M4 module**

From BR7M4, we extracted DEGs for drug discovery. The DEGs in the BR7M4 module were obtained by analyzing the data from GPL96 (Table S6). In total, 9 up-regulated genes and 163 down-regulated genes in BR7M4 were used for drug discovery by CMAP. In terms of the CMAP output, the connectivity score tending to -1 indicates strong restoration of expression of PD dysregulated genes. As shown in Table S9, using the connectivity score -0.8 as a cutoff, we selected 14 drug candidates for PD therapy. Information on these drug candidates is provided in Table S9. Of these, sodium phenylbutyrate (SP) and meclofenoxate hydrochloride (MH) are capable of passing through the blood brain barrier, and so their effectiveness for PD was further investigated.

**Predicting drug targets for MH and SP**

Further analyses were carried out to examine the potential target proteins for MH or SP by using DStruBTarget[2] to predict those proteins that could directly bind to MH and SP. The drug structure similarity between MH and SP is 0.143 as measured by 2D Tanimoto similarity, indicating that these two drugs are largely dissimilar in their structures when the suggested drug similarity cutoff was set to 0.4[3, 4]. The drug structures of MH and SP were input into DStruBTarget that predicted the top 10 candidate proteins potentially bound to them (Table S10). Function enrichment analysis by ClusterProfiler[5] indicated that the top 10 proteins predicted to bind MH were enriched in neuroactive ligand-receptor interactions and neurotransmitter receptor activity functions (*P* =1.27$\times$10^-7^), whereas the top 10 proteins predicted to bind with SP were enriched in inflammation-related functions (*P* < 0.05) (Table S11), further suggesting differential mechanisms of action for MH and SP. Among the potential targets of MH shown in Table S10, CNR1 (Cannabinoid receptor 1), one protein target of MH, was a G-protein previously shown to induce neurite outgrowth in Neuro2A cells through activation of Src kinase[6, 7].

**MH and SP can improve glucose metabolism in the brains of ROT-induced PD mice**

In this study, 86 C57B/L mice were randomly divided into 6 groups viz. DMSO (NC), ROT, SP, MH, ROT+SP and ROT+MH groups. In total, 15 mice were in the NC group, 13 mice were in the SP group, 16 mice were in the MH group and 12 mice were in the ROT+MH group (Table S8). From these mice, 28 were randomly selected to test glucose metabolism in their brains by employing neuro-imaging with [^18^F]-fluorodeoxyglucose positron emission tomography (^18^F- FDG PET): 5 mice from the NC group, 5 mice from the ROT group, 4 mice from the SP group, 6 mice from the MH group, 5 mice from the ROT+SP group and 3 mice from the ROT+MH group. Neuro-imaging with [^18^F]-fluorodeoxyglucose positron emission tomography (^18^F- FDG PET) can capture synaptic dysfunction *in vivo*. The radiotracer ^18^F-FDG provides an index for the cerebral metabolic rate of glucose, which is strongly associated with neuronal activity and synaptic integrity[8, 9].

**Impact of SP on the homeostasis of mitochondrial marker proteins**

The level of α-synuclein exhibited by mice in the ROT group increased from 1.06 to 1.35 (*P* = 0.0035, *P* < 0.01) as compared to the NC group, whilst expression of α-synuclein decreased from 1.35 to 0.90 in the ROT + SP group (P = 0.028) compared to the ROT group (Fig. 6F). The NeuN level of mice in the ROT group decreased from 0.99 to 0.50 (P = 0.025) as compared to the NC group, whereas ROT+SP groups displayed significant increases (from 0.50 to 0.89) in NeuN expression (P = 0.0078, *P < 0.01*) as compared to the ROT group (Fig. 6G).

**Impact of MH on the genes involved in pathways pertaining to mitochondrial function and postsynaptic density**

We used qRT-PCR to test a set of genes (*Ndufa12, Cox6b1, Atp5k, Src, Ndufb10* and *Dlgap3*) in the Uni-MH-ROT that are known to be involved in the pathways pertaining to mitochondrial function and postsynaptic density (Fig. S6A). Ndufa12 increased ~ 1.4 fold in the ROT group (*P* < 0.001) compared to the NC group. Upon MH treatment, the expression of *Ndufa12* was reduced significantly (*P* < 0.01) by comparison with the ROT-induced group, to a level comparable to the NC group (Fig. 7A). The level of *Cox6b1* expression was up-regulated ~1.2 fold in the ROT group (P<0.001) but was restored to a normal level upon MH treatment (*P* < 0.001) (Fig. 7B). Similarly, the levles of *Atp5k* (~1.2 fold, *P < 0.01*) (Fig. 7C) and *Src* (~1.5 fold, *P* < 0.05) (Fig. 7D) expression were up-regulated after ROT treatment, but normal after MH treatment.

To better understand how MH might impact synaptic function in the PD mouse model, qRT-PCR was performed to assess the drug’s effect on the expression of several genes (*Chrna4, Syt11, Cdh8* and *Sncg*) that are known to play a role in synaptic pathways[10-14] in the NC, ROT, MH or MH+ROT groups. As shown in Fig. 7E, the *Chrna4* expression level increased ~1.4 fold in the ROT group (*P* =4.0$\times10$^-2^) compared to that of the NC group whereas *Chrna4* expression in the MH+ROT group was restored to near normal, significantly (*P* =3.2$\times10$^-2^) lower than in the ROT group. The expression of the *Syt11* gene increased ~1.2 fold in the ROT group (*P* =2.0$\times10$^-1^) but was restored almost back to that of the NC group upon MH treatment (Fig. 7F). Similarly, the *Cdh8* gene (~2-fold, *P* =1.4$\times10$^-2^) (Fig. 7G) was down-regulated by MH compared to the ROT group. The expression of the *Sncg* gene increased significantly after ROT treatment compared with the NC group (*P* =2.1$\times10$^-2^), whereas MH treatment (MH+ROT group) restored expression to a level close to that of the NC group (*P* =7.4$\times10$^-1^) although no significant difference was observed between the ROT and MH+ROT groups (Fig. 7H). These data clearly indicate that MH is capable of modulating the expression of these genes, and contribute to protect synaptic function in PD.

We also used qRT-PCR to test a set of genes (*Ndufb10, Dlgap3, APBA1, GLRA1* and *NEFH*) in the Uni-MH-ROT that are known to be involved in the pathways pertaining to mitochondrial function and postsynaptic density (Fig. S6A). However, there was no significant difference in the expression of these genes(Fig. S8A-E).

**Supplementary Tables**

**Table S1. Number of genes and PD-associated gene or SNP enrichment (respectively by chi-square test and by sLDSC enrichment analysis) in each consensus co-expressed module (CCM). The P-values were transformed into –log­_10_(P-values)**

|  |  | **PD-genes (DisGeNet)** | | | | | **PD-SNPs (Chi-square)** | | | | **PD-SNPs(sLDSC)** | | | | |
| --- | --- | --- | --- | --- | --- | --- | --- | --- | --- | --- | --- | --- | --- | --- | --- |
| **ID** | **# genes** | **G1^a^** | **G2^b^** | **G3^c^** | **G4^d^** | **G5^e^** | **S1^f^** | **S2^g^** | **S3^h^** | **S4^i^** | **S1** | **S2** | **S3** | **S4** | **S5^j^** |
| M0 | 1121 | 0.509 | 0.173 | 0.040 | 0.000 | 0.635 | 0.000 | 0.000 | 0.067 | 1.483 | 0.000 | 0.785 | 0.130 | 0.000 | 0.110 |
| M1 | 7990 | 0.019 | 0.180 | 0.246 | 0.740 | 0.001 | 0.117 | 0.400 | 0.207 | 1.291 | 0.366 | 0.166 | 0.237 | 2.363 | 8.463 |
| M2 | 7113 | 0.000 | 0.000 | 0.000 | 0.006 | 0.129 | 0.152 | 0.011 | 0.000 | 0.005 | 0.439 | 0.000 | 0.000 | 0.256 | 1.307 |
| M3 | 2791 | 5.030 | 8.208 | 2.220 | 1.075 | 0.733 | 0.000 | 0.170 | 4.389 | 0.306 | 0.055 | 0.483 | 1.812 | 0.203 | 1.924 |
| M4 | 1434 | 0.535 | 1.065 | 2.342 | 0.000 | 0.483 | 0.450 | 0.359 | 0.943 | 0.048 | 0.000 | 0.796 | 0.363 | 1.220 | 0.287 |
| M5 | 292 | 3.024 | 3.558 | 2.036 | 0.474 | 0.000 | 1.660 | 0.000 | 0.350 | 0.511 | 0.000 | 0.000 | 0.000 | 0.000 | 0.357 |
| M6 | 295 | 0.572 | 1.582 | 1.413 | 0.000 | 0.000 | 0.682 | 1.003 | 0.348 | 0.145 | 1.361 | 0.670 | 0.000 | 0.000 | 1.108 |
| M7 | 234 | 0.057 | 0.161 | 1.303 | 0.000 | 0.000 | 0.000 | 0.000 | 0.426 | 0.210 | 0.117 | 0.027 | 0.041 | 0.000 | 0.426 |
| M8 | 227 | 2.907 | 4.575 | 3.211 | 0.000 | 0.000 | 0.000 | 1.116 | 0.963 | 1.631 | 0.279 | 0.188 | 0.000 | 0.000 | 0.815 |
| M9 | 180 | 3.441 | 4.353 | 1.413 | 0.652 | 2.032 | 0.000 | 0.000 | 0.585 | 0.040 | 0.190 | 0.550 | 0.000 | 0.000 | 0.179 |
| M10 | 167 | 0.790 | 0.234 | 0.183 | 1.648 | 0.000 | 0.917 | 0.000 | 0.243 | 0.214 | 0.000 | 0.198 | 0.000 | 0.000 | 0.000 |
| M11 | 122 | 0.000 | 0.002 | 0.000 | 0.000 | 0.000 | 0.975 | 0.000 | 0.282 | 0.078 | 0.189 | 0.000 | 0.000 | 0.000 | 0.000 |
| M12 | 113 | 0.020 | 1.662 | 0.049 | 0.000 | 0.000 | 0.000 | 0.000 | 0.323 | 0.333 | 0.000 | 0.000 | 0.000 | 0.000 | 0.234 |
| M13 | 44 | 2.975 | 2.305 | 1.239 | 1.225 | 0.000 | 1.313 | 0.000 | 0.000 | 0.000 | 0.385 | 0.148 | 1.190 | 1.252 | 0.290 |
| M14 | 59 | 0.097 | 0.304 | 0.063 | 0.000 | 0.000 | 0.000 | 0.000 | 0.567 | 0.000 | 0.182 | 0.000 | 0.000 | 0.000 | 0.000 |
| M15 | 58 | 1.714 | 0.544 | 1.364 | 1.109 | 0.000 | 0.000 | 0.000 | 0.000 | 0.000 | 0.177 | 0.179 | 0.000 | 0.097 | 0.588 |
| M16 | 46 | 2.870 | 4.804 | 4.764 | 1.206 | 1.433 | 0.000 | 0.000 | 0.000 | 0.000 | 0.205 | 0.014 | 0.000 | 0.000 | 0.991 |
| M17 | 36 | 0.205 | 1.166 | 0.471 | 0.000 | 0.000 | 0.000 | 0.000 | 0.000 | 0.000 | 0.000 | 1.265 | 0.000 | 0.176 | 0.000 |
| M18 | 19 | 0.000 | 0.000 | 0.000 | 0.000 | 0.000 | 0.000 | 0.000 | 0.000 | 0.511 | 0.040 | 0.259 | 0.000 | 0.000 | 0.165 |

^a^AlteredExpression; ^b^Biomarker; ^c^GeneticVariation; ^d^PosttranslationalModification; ^e^Therapeutic; ^f^GCST007780; ^g^GCST009373; ^h^GCST010765; ^i^Lancet2019; ^j^ GCST009374.

**Table S2. Similarities of 10 brain regions evaluated by the number of overlapping genes (Chi-square test) between one pair of modules from different brain regions**

|  | **CEC^a^** | **FRC^b^** | **HIP^c^** | **MED^d^** | **OCC^e^** | **PUT^f^** | **SUN^g^** | **TEC^h^** | **THA^i^** | **WHM^g^** |
| --- | --- | --- | --- | --- | --- | --- | --- | --- | --- | --- |
| CEC | 3.68 | 21.93 | 11.70 | 8.60 | 32.42 | 19.91 | 11.61 | 36.71 | 14.09 | 14.01 |
| FRC | 21.93 | 3.52 | 39.60 | 25.48 | 40.26 | 15.31 | 31.78 | 46.01 | 27.12 | 42.30 |
| HIP | 11.70 | 39.60 | 4.83 | 5.98 | 20.14 | 11.19 | 13.35 | 22.57 | 9.90 | 9.95 |
| MED | 8.60 | 25.48 | 5.98 | 3.93 | 14.21 | 8.36 | 11.82 | 15.40 | 8.62 | 8.99 |
| OCC | 32.42 | 40.26 | 20.14 | 14.21 | 3.87 | 19.28 | 11.59 | 16.66 | 21.26 | 18.86 |
| PUT | 19.91 | 15.31 | 11.19 | 8.36 | 19.28 | 3.49 | 16.29 | 31.85 | 14.52 | 16.31 |
| SUN | 11.61 | 31.78 | 13.35 | 11.82 | 11.59 | 16.29 | 3.46 | 55.99 | 15.15 | 16.05 |
| TEC | 36.71 | 46.01 | 22.57 | 15.40 | 16.66 | 31.85 | 55.99 | 5.76 | 26.78 | 55.97 |
| THA | 14.09 | 27.12 | 9.90 | 8.62 | 21.26 | 14.52 | 15.15 | 26.78 | 3.44 | 11.91 |
| WHM | 14.01 | 42.30 | 9.95 | 8.99 | 18.86 | 16.31 | 16.05 | 55.97 | 11.91 | 3.46 |

^a^Cerebellar cortex; ^b^ Frontal cortex; ^c^ Hippocampus; ^c^ Medulla; ^d^ Occipital cortex; ^e^ Putamen; ^f^ Substantia nigra; ^g^ Temporal cortex; ^h^ Thalamus; ^i^ White matter. The number in each cell is -log (FDR) of Chi-square test.

**Table S3. PD-associated genes collated by DisGeNet database**

| **Association type** | **Number of genes** |
| --- | --- |
| AlteredExpression | 1518 |
| Biomarker | 7024 |
| CausalMutation | 1 |
| GeneticVariation | 4577 |
| PosttranslationalModification | 52 |
| Therapeutic | 23 |

**Table S4. Information on each GWAS summary file for the extraction of PD-associated SNPs**

| **GWAS Summary** | **Publishing Time** | **PubMed ID** | **Sample Size** | **Number of PD-associated SNPs** | **Number of Exons within 10kb** |
| --- | --- | --- | --- | --- | --- |
| GCST010765 | 2020-06-23 | 32589924 | 28,568 | 399 | 131 |
| GCST009373 | 2019-11-22 | 31755958 | 10,107 | 61 | 9 |
| GCST009374 | 2019-11-22 | 31755958 | 1,353 | 0 | 0 |
| GCST007780 | 2019-04-07 | 30957308 | 1,345 | 139 | 20 |
| Lan2019 | 2019-12-01 | 31701892 | 56,306 | 5806 | 319 |

**Table S5. Number of genes and PD-associated gene or SNP enrichment (respectively by chi-square test and by sLDSC enrichment analysis) in each of the four brain region specific co-expressed modules (SCM). The P-values were transformed into –log_10_(P-values)**

|  |  |  | **PD-genes (DisGeNet)** | | | | | **PD-SNPs (Chi-square)** | | | | **PD-SNPs(sLDSC)** | | | | |
| --- | --- | --- | --- | --- | --- | --- | --- | --- | --- | --- | --- | --- | --- | --- | --- | --- |
| **ID** | **Brain region** | **# genes** | **G1^a^** | **G2^b^** | **G3^c^** | **G4^d^** | **G5^e^** | **S1^f^** | **S2^g^** | **S3^h^** | **S4^i^** | **S1** | **S2** | **S3** | **S4** | **S5^j^** |
| BR7M4 | Substantia nigra | 399 | 3.666 | 5.810 | 10.298 | 4.799 | 1.383 | 1.389 | 0.866 | 1.703 | 0.386 | 1.614 | 0.582 | 0.000 | 0.775 | 0.842 |
| BR9M3 | Thalamus | 1934 | 4.009 | 4.208 | 1.148 | 1.827 | 0.686 | 0.090 | 0.276 | 2.745 | 0.233 | 0.038 | 1.492 | 1.585 | 0.282 | 1.301 |
| BR6M3 | Putamen | 2142 | 6.009 | 5.265 | 1.790 | 2.131 | 1.046 | 0.266 | 0.726 | 4.535 | 0.580 | 0.078 | 0.714 | 1.409 | 0.243 | 2.060 |
| BR3M2 | Hippocampus | 2296 | 4.307 | 7.519 | 1.476 | 1.955 | 0.959 | 0.000 | 0.000 | 4.548 | 0.611 | 0.235 | 0.241 | 0.724 | 0.033 | 2.208 |

^a^AlteredExpression; ^b^Biomarker; ^c^GeneticVariation; ^d^PosttranslationalModification; ^e^Therapeutic; ^f^GCST007780; ^g^GCST009373; ^h^GCST010765; ^i^Lancet2019; ^j^ GCST009374.

**Table S6. GEO datasets for detecting differentially expressed genes (DEGs) between PD and controls**

| **Dataset** | **Brain region** | **Platform** | **Sample size of controls** | **Sample size of PD** |
| --- | --- | --- | --- | --- |
| GSE8397 | Lateral substantia nigra | GPL96 | 7 | 9 |
|  | Medial substantia nigra |  | 8 | 15 |
|  | Total |  | 15 | 24 |
|  | Lateral substantia nigra | GPL97 | 7 | 9 |
|  | Medial substantia nigra |  | 8 | 15 |
|  | Total |  | 15 | 24 |

**Table S7. GEO Datasets used for the module conservation test**

| **Dataset ID** | **Country of samples** | **Tissue** | **Groups** | **Sample size** |
| --- | --- | --- | --- | --- |
| GSE131617 | Japan | Entorhinal cortex | Braak NFT stages of 0 | 13 |
|  |  |  | Braak NFT stages of I–II | 20 |
|  |  |  | Braak NFT stages of III–IV | 19 |
|  |  |  | Braak NFT stages of V–VI | 19 |
|  |  | Temporal cortex | Braak NFT stages of 0 | 13 |
|  |  |  | Braak NFT stages of I–II | 20 |
|  |  |  | Braak NFT stages of III–IV | 19 |
|  |  |  | Braak NFT stages of V–VI | 19 |
|  |  | Frontal cortex | Braak NFT stages of 0 | 13 |
|  |  |  | Braak NFT stages of I–II | 20 |
|  |  |  | Braak NFT stages of III–IV | 19 |
|  |  |  | Braak NFT stages of V–VI | 19 |
| GSE23290 | Spain | Putamen | Controls | 5 |
|  |  |  | PD | 8 |
| GSE34516 | Spain | Locus coeruleus | Controls | 4 |
|  |  |  | PD | 6 |
| GSE51922 | Spain | IPSC-induced dopaminergic neurons | Controls | 4 |
|  |  |  | PD | 6 |
| GSE18838 | Miami, USA | Peripheral blood | Controls | 11 |
|  |  |  | PD | 17 |
| GSE34862 | UK | Substantia Nigra | Controls | 57 |

**Table S8. Number of mice used in each experiment**

| **Group** | **PET-CT & RNAseq** | **IHC** | **Behavior & mitochondria** | **Total** | **Weight**  **(before treatment)** | **Weight**  **(after treatment)** |
| --- | --- | --- | --- | --- | --- | --- |
| NC | 5 | 4 | 6 | 15 | 26.75±1.15 | 26.99±0.95 |
| SP | 4 | 3 | 6 | 13 | 26.90±1.20 | 26.94±0.59 |
| ROT+SP | 5 | 4 | 6 | 15 | 26.31±0.86 | 26.03±1.03 |
| MH | 6 | 4 | 6 | 16 | 26.47±1.78 | 26.44±1.00 |
| ROT+MH | 3 | 3 | 6 | 12 | 25.95±1.05 | 25.95±0.74 |
| ROT | 5 | 4 | 6 | 15 | 26.85±1.17 | 26.90±0.99 |
| Total | 28 | 22 | 36 | 86 |  |  |

**Table S9. Drug candidates predicted to restore the DEGs in the BR7M4 module.**

| **Rank** | **Drug name** | **Connectivity score** | **Cell type** | **Dose** (µM) |
| --- | --- | --- | --- | --- |
| 1 | Bacampicillin | -1 | MCF7 | 8 |
| 2 | Sodium phenylbutyrate | -0.963 | MCF7 | 1000 |
| 3 | Cyanocobalamin | -0.877 | MCF7 | 3 |
| 4 | Metaraminol | -0.85 | MCF7 | 9 |
| 5 | Primaquine | -0.846 | HL60 | 9 |
| 6 | SC-58125 | -0.836 | HL60 | 10 |
| 7 | Dexamethasone | -0.835 | ssMCF7 | 1 |
| 8 | Levomepromazine | -0.825 | PC3 | 9 |
| 9 | Carteolol | -0.82 | MCF7 | 12 |
| 10 | Rofecoxib | -0.819 | MCF7 | 10 |
| 11 | Folic acid | -0.816 | PC3 | 9 |
| 12 | Meclofenoxate | -0.814 | MCF7 | 14 |
| 13 | Cefixime | -0.806 | MCF7 | 9 |
| 14 | Clidinium bromide | -0.806 | PC3 | 9 |

**Table S10.** **Predicted human protein targets binding with MH and SP by DStruBTarget**

|  | **UniProt** | **Gene symbol** | **Score** |
| --- | --- | --- | --- |
| **MH** | P07148 | *FABP1* | 0.618548 |
|  | Q99720 | *SIGMAR1* | 0.608120 |
|  | P37231 | *PPARG* | 0.604557 |
|  | P21917 | *DRD4* | 0.575678 |
|  | P21554 | *CNR1* | 0.557235 |
|  | P25100 | *ADRA1D* | 0.542949 |
|  | P34972 | *CNR2* | 0.536092 |
|  | P08908 | *HTR1A* | 0.528512 |
|  | P10323 | *ACR* | 0.524715 |
|  | P14324 | *FDPS* | 0.524260 |
| **SP** | P15085 | *CPA1* | 0.746952 |
|  | Q9H4A4 | *RNPEP* | 0.712588 |
|  | P55211 | *CASP9* | 0.709387 |
|  | P42574 | *CASP3* | 0.707630 |
|  | Q16719 | *KYNU* | 0.707037 |
|  | P29466 | *CASP1* | 0.702633 |
|  | Q9UJM8 | *HAO1* | 0.655972 |
|  | P27695 | *APEX1* | 0.637906 |
|  | Q13133 | *NR1H3* | 0.629362 |
|  | P28838 | *LAP3* | 0.628151 |

**Table S11. KEGG enrichment analysis of predicted protein targets respectively for MH and SP**

| **Drug** | **KEGG term** | **Adjusted**  ***P*-value** | **Binding targets** |
| --- | --- | --- | --- |
| **MH** | Neuroactive ligand-receptor interaction | 1.27×10^-7^ | DRD4,PTGER2,HTR1A,CNR1HTR1B,CNR2,ADRA1D,HTR2A |
|  | Serotonergic synapse | 1.40×10^-2^ | HTR1A,HTR1B,HTR2A |
| **SP** | Legionellosis | 4.00×10^-3^ | CASP3,CASP9,CASP1 |
|  | Apoptosis – multiple species | 4.07×10^-2^ | CASP3,CASP9 |
|  | Pathogenic *Escherichia coli* infection | 4.07×10^-2^ | CASP3,CASP9,CASP1 |
|  | Hepatitis C | 4.07×10^-2^ | CASP3,CASP9,NR1H3 |
|  | Influenza A | 4.07×10^-2^ | CASP3,CASP9,CASP1 |

**Table S12.** **Metabolites expressed significantly differently between MH+ROT and ROT groups**

| **Name** | **OPLS VIP** | **t-test *P*-value** |
| --- | --- | --- |
| 17a-Ethynylestradiol | 1.391 | 2.77×10^-4***^ |
| L-Indospicine | 1.107 | 4.84×10^-4***^ |
| 2-{(3S)-1-[4-(Trifluoromethyl)benzyl]-3-pyrrolidinyl}-1,3-benzoxazole | 1.574 | 5.35×10^-3**^ |
| Cotinine | 1.481 | 2.11×10^-2*^ |
| Cypridinaluciferin | 1.220 | 3.59×10^-2*^ |
| 5-HETE | 1.250 | 3.9×10^-2*^ |
| D-(+)-Maltose | 1.034 | 4.78×10^-2*^ |
| Oxprenolol | 1.562 | 4.99×10^-2*^ |
| trans-Cinnamoyl beta-D-glucoside | 1.151 | 6.28×10^-2^ |
| L-Tryptophanamide | 1.097 | 6.54×10^-2^ |
| Zearalanol | 1.354 | 6.72×10^-2^ |
| Estrone | 1.039 | 6.87×10^-2^ |
| 17beta-hydroxy-5beta-androstan-3-one | 1.084 | 8.08×10^-2^ |
| 6,9-dioxo-11R,15S-dihydroxy-13E-prostenoic acid | 1.113 | 8.20×10^-2^ |
| 2-Heptyl-4-hydroxyquinoline-N-oxide | 1.321 | 8.56×10^-2^ |
| L-Tyrosine | 1.246 | 8.89×10^-2^ |
| Dihydrothymine | 1.195 | 8.94×10^-2^ |
| DL-Carnitine | 1.178 | 9.06×10^-2^ |
| 6-Methoxyquinoline | 1.174 | 9.20×10^-2^ |
| 5,5-dimethyl-2-{[(2-phenylacetyl)amino]methyl}-1,3-thiazolane-4-carboxylic acid | 1.223 | 9.37×10^-2^ |
| N-Methylpyrrolidone | 1.153 | 9.62×10^-2^ |
| Oleic acid-biotin | 1.179 | 9.83×10^-2^ |

OPLS VIP: Variable influence on projection of Orthogonal projections to latent structures

**Table S13. The primer sequences.**

| **Gene** | **The primer sequences(5'to3')** |
| --- | --- |
| Ndufa12 F | CGGCAAAAACACATTCTGGG |
| Ndufa12 R | GGAGTGGCAGACACATTGAA |
| Cox6b1 F | AACTGCCCCCTTTGACAGC |
| Cox6b1 R | CACACACGGAGACATCACC |
| Atp5k F | GTTCAGGTCTCTCCACTCAT |
| Atp5k R | CCGTTTCAACTCATCTAGTCT |
| Src F | TCACTAGACGGGAATCAGAG |
| Src R | GCTTGCGGATCTTGTAGTGT |
| CHRNA4 F | ACTACGAGAATGTCACCTCC |
| CHRNA4 R | AAGAAGGTGACGTCGATGCT |
| SNCA F | CACTGGCTTTGTCAAGAAGGACC |
| SNCA R | CATAAGCCTCACTGCCAGGATC |
| SYT11 F | TTAGCATCTACCCAGAGACC |
| SYT11 R | CCATACAAGATGCAGGGCTA |
| CDH8 F | TTGACCGAGAGGAAAAGGCT |
| CDH8 R | GGACATCTCTGGAACAGTAG |
| GAPDH F | AGGTCGGTGTGAACGGATTTG |
| GAPDH R | TGTAGACCATGTAGTTGAGGTCA |
| Ndufb10 F | ACCTACTACTACCACCGACA |
| Ndufb10 R | CCTGGTCAACTTTGAAGTCC |
| Dlgap3 F | ATGGACAGTCAGTCAAGCGA |
| Dlgap3 R | TGATAAGTCCTGGCTTTGGC |
| APBA1 F | AAGCAGAGCATGAGTTCGCA |
| APBA1 R | CTCCTTGATGTCCTTGATGG |
| GLRA1 F | ACCAAGCACTACAACACAGG |
| GLRA1 R | ATGGTGAGCACTGTGGTGAT |
| NEFH F | GCCAAAGTGAACACAGATGC |
| NEFH R | ACTGTCCAGCTGCTGAATAG |

**Table S14.** **Gradient elution ratio**

| **Time (min)** | **A (%)** | **B（%）** |
| --- | --- | --- |
| 0 | 98 | 2 |
| 0.5 | 98 | 2 |
| 12 | 50 | 50 |
| 14 | 98 | 2 |
| 16 | 98 | 2 |
| 16.1 | 2 | 98 |
| 23 | 2 | 98 |

Phase A: 95:5 (acetonitrile:water) with 10 mM ammonium formate, 0.1% formic acid; Phase B: 50:50 (acetonitrile:water) with 10 mM ammonium formate, 0.1% formic acid

**Supplementary Figures**

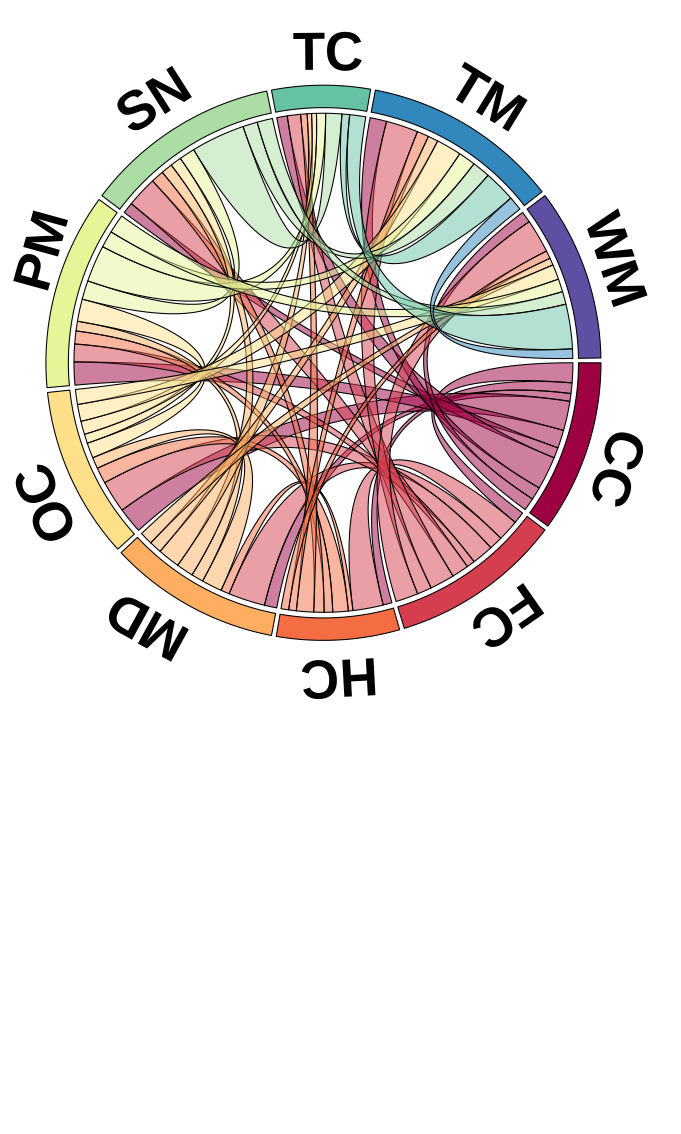

**Fig. S1. Gene co-expression modules specifically expressed in one of the ten regions of healthy brains.** The length of the rectangle represents the number of co-expression modules specifically expressed in the corresponding brain region. The width of the link between two modules represents the significance (single-tailed hyper-geometric test) of the overlapping genes between two modules as compared to the genes in the other modules. The wider the link, the more significant the overlap.

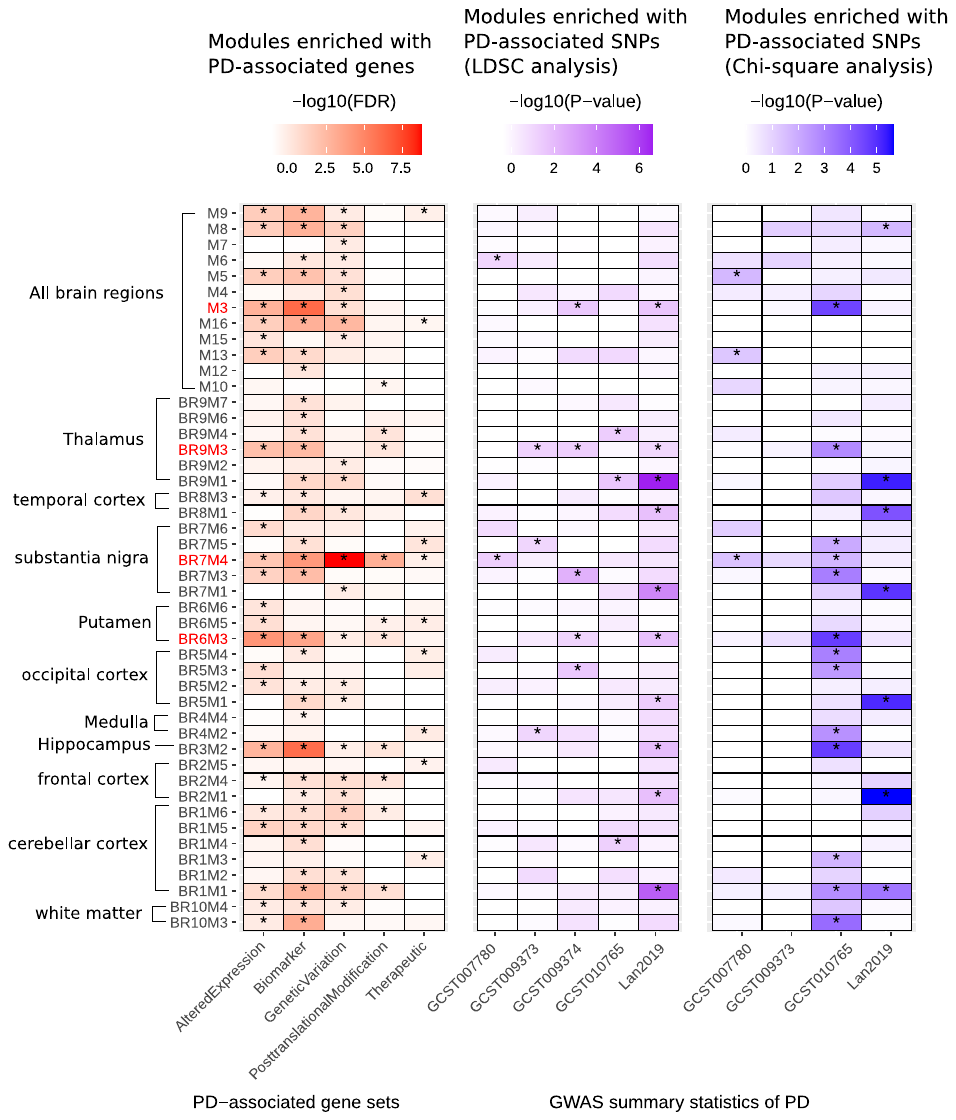

**Fig. S2.** **Enrichment of PD-associated genes or SNPs in SCM and CCM modules.**

“*” indicated *P* <0.05. *P*-values were calculated by single-tailed Fisher’s exact test. SCM: gene co-expression modules specifically expressed in certain brain regions;

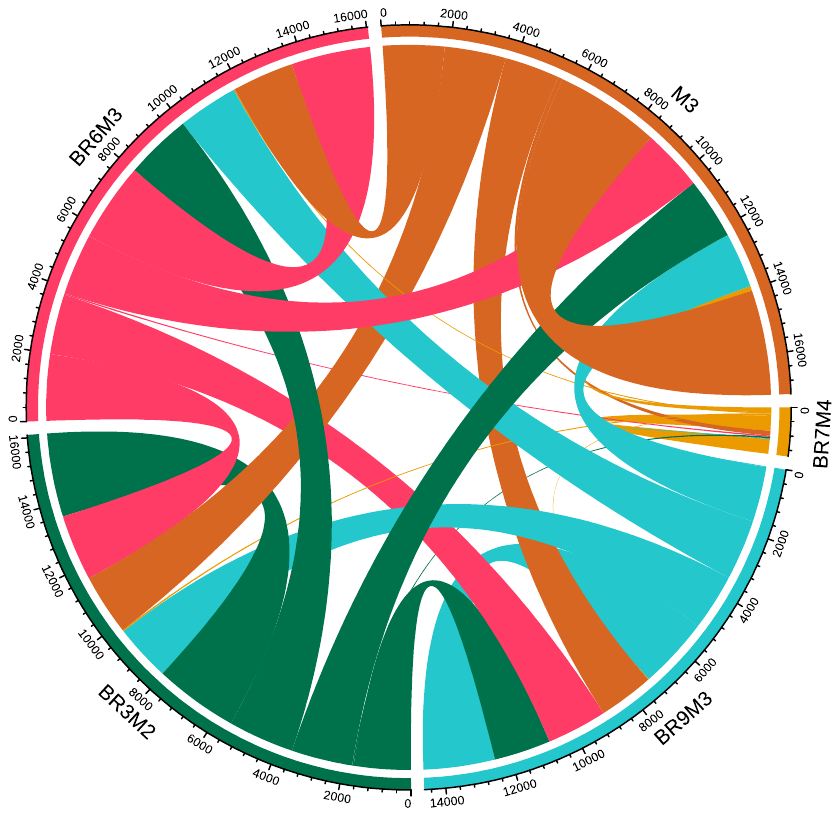

**Fig. S3. Overlapping genes among modules M3, BR9M3, BR7M4, BR6M3 and BR3M2.**
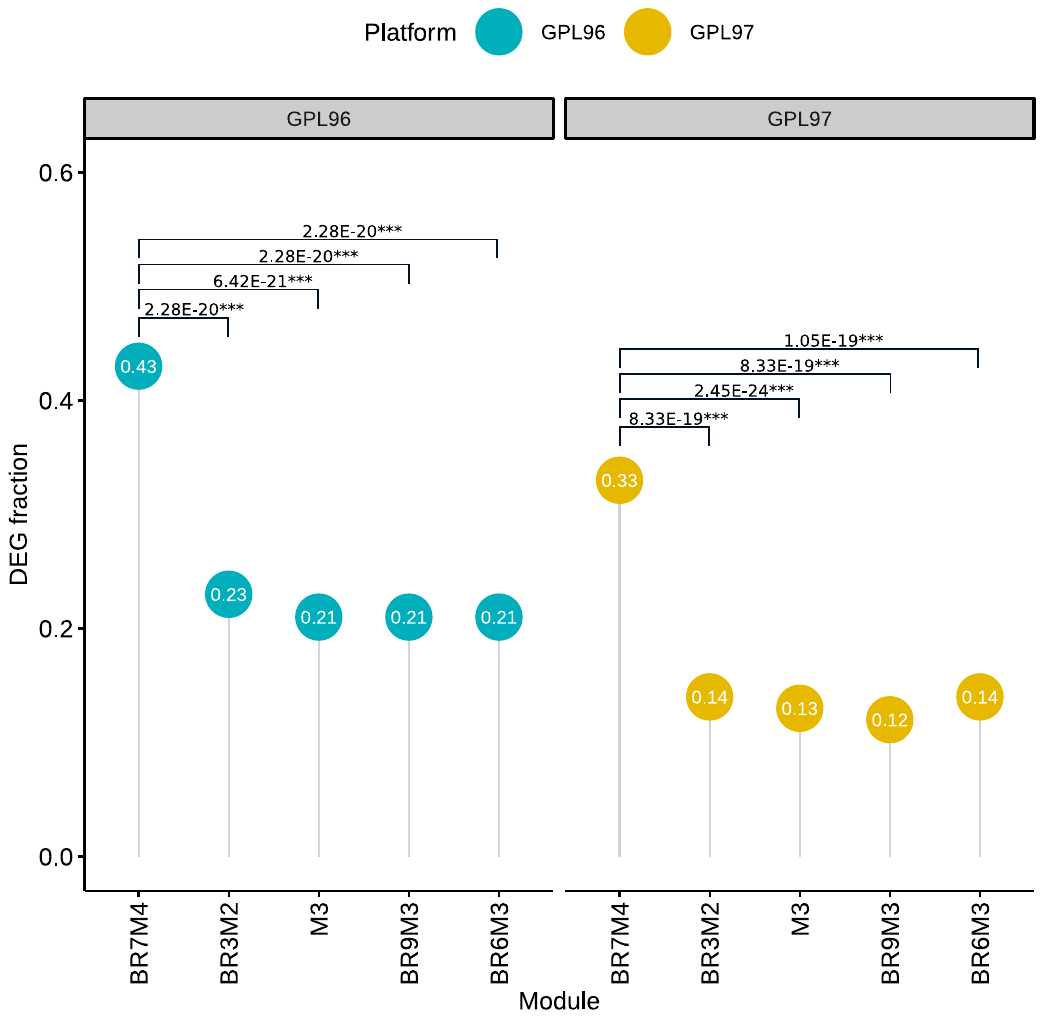

**Fig. S4. The proportions of genes expressed significantly differently between PD patients and controls (DEGs) in PD-associated modules including BR7M4, BR3M2, BR9M3, BR6M3 and M3.**

The DEGs were genes obtained from two platforms (GPL96 and GPL97, respectively). “***” indicates single-tailed Chi-square test adjusted *P* < 0.001.

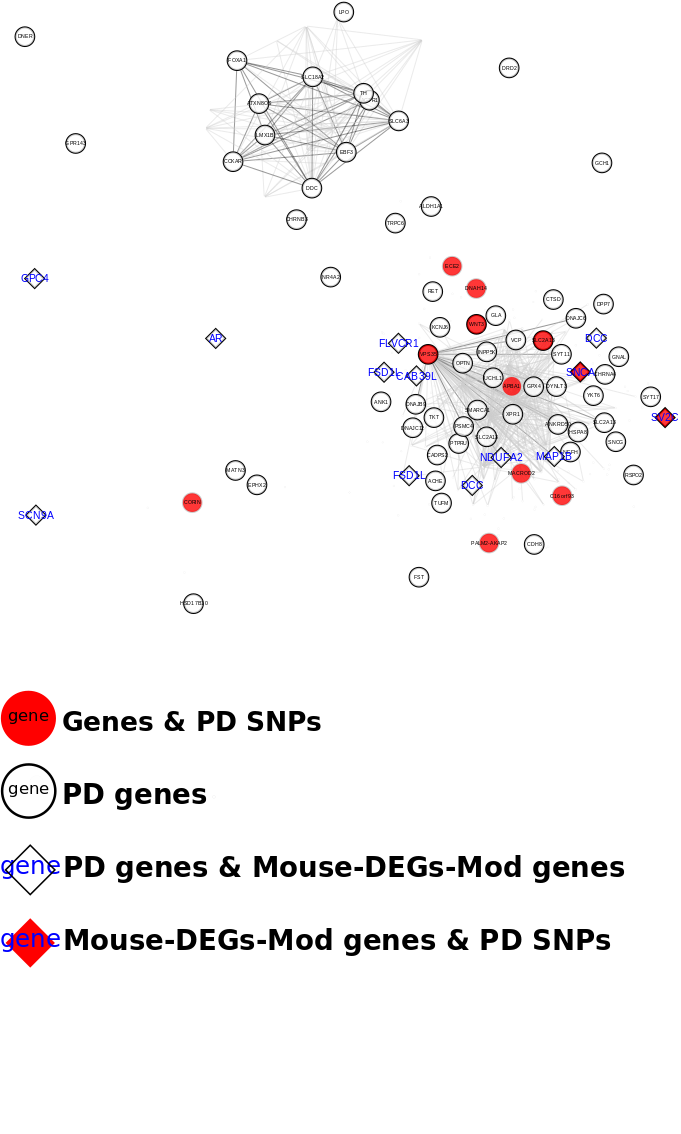

**Fig. S5. Co-expression network of BR7M4.** PD-associated genes collated by DisGeNet are highlighted by black circles. Genes located within 10kb of the PD-associated SNPs (by Chi-square test) are colored in red. The genes (Mouse-DEGs) expressed significantly differently between the ROT and NC groups are denoted by diamonds with light green labels. Other genes in BR7M4 are colored light grey. The expression co-relationship between each pair of genes is represented by edge length, and the co-relationship between PD-associated genes is indicated by black lines.

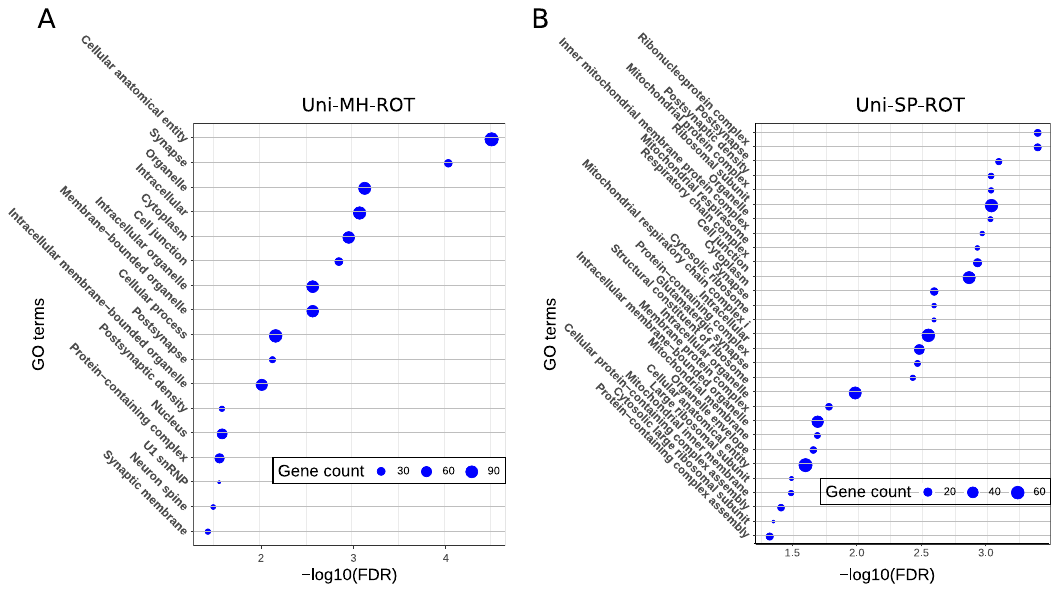

**Fig. S6. GO function enrichment analysis of genes expressed significantly differently between the ROT group and SP-treated or MH-treated cells.**

**(A)** GO functions enriched by protein-protein interactions in the Uni-MH-ROT set. **(B)** GO functions enriched by protein-protein interactions in the Uni-SP-ROT set.

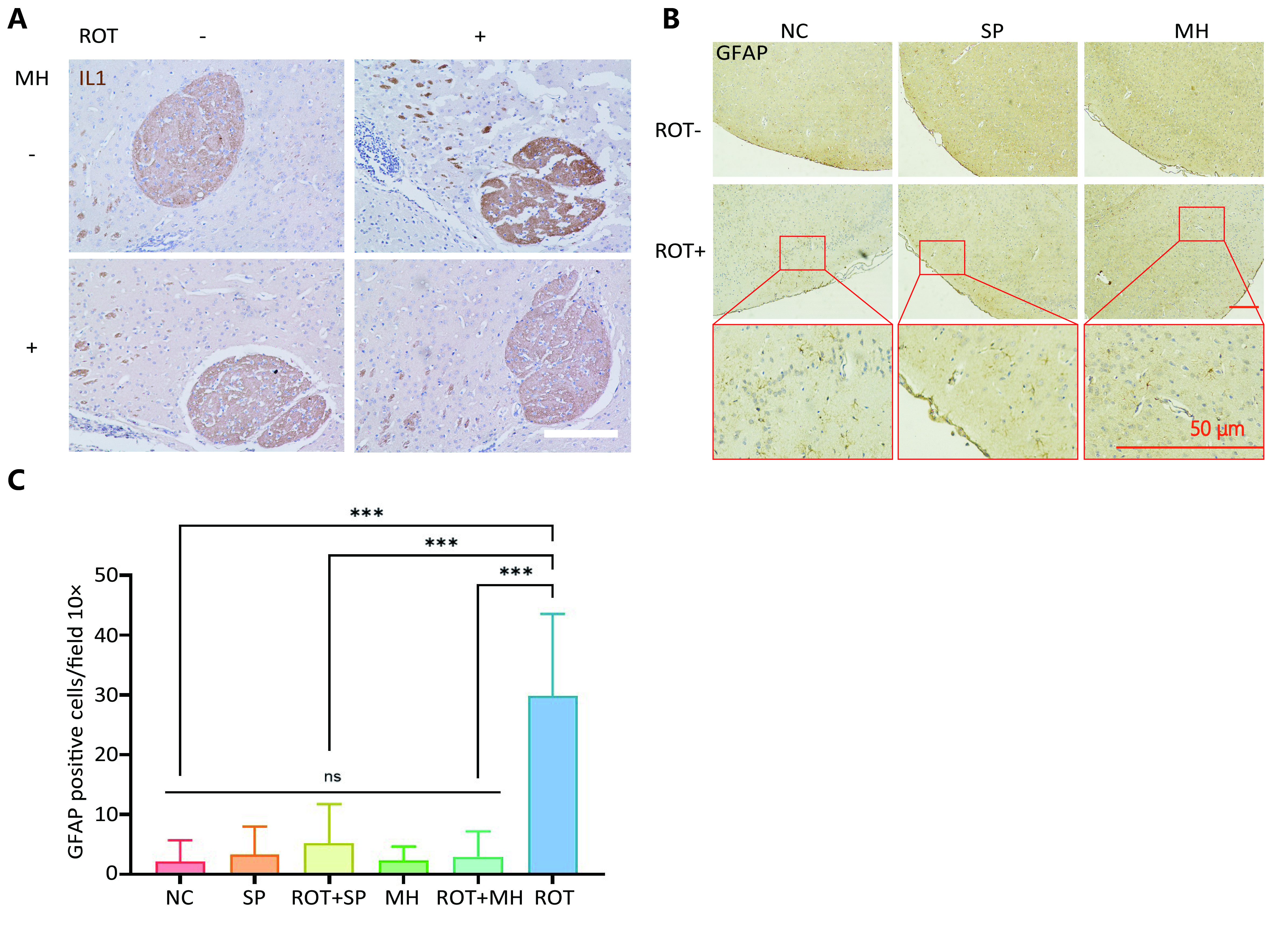

**Fig. S7. MH prevents ROT-induced neurodegeneration.**

**(A)** Immunohistochemistry was used to compare the content of IL1-positive cells in brain slices from the NC, ROT, MH and ROT+MH groups, and the staining between each group (not less than 3 in each group). It can be seen that the staining of IL1-positive cells in the ROT group was significantly deeper than that of the NC group, whereas the staining of the MH group was similar to that of the NC. The staining in the ROT+MH group indicated that MH treatment can effectively reverse the increase in number of IL1-positive cells caused by ROT. **(B)** The proportions of GFAP+ cells were markedly increased in the striatum of the ROT group (29.81±13.75) compared to that in the NC group (2.16±3.52) (*P* = 2.95$\times$10^-11^). By contrast, the numbers of GFAP^+^ cells in the striatums of the ROT+SP and ROT+MH groups were significantly lower than that of the ROT group [SP: 5.23 ± 6.51 (*P* = 1.64$\times$10^-11^) and MH: 2.96 ± 4.22 (*P* = 1.67$\times$10^-11^)]. (C) Little or no difference was observed in terms of the numbers of GFAP+ cells between the ROT+SP and ROT+MH groups. A:Scale bar = 10 *μm,* B:Scale bar = 50 *μm*.

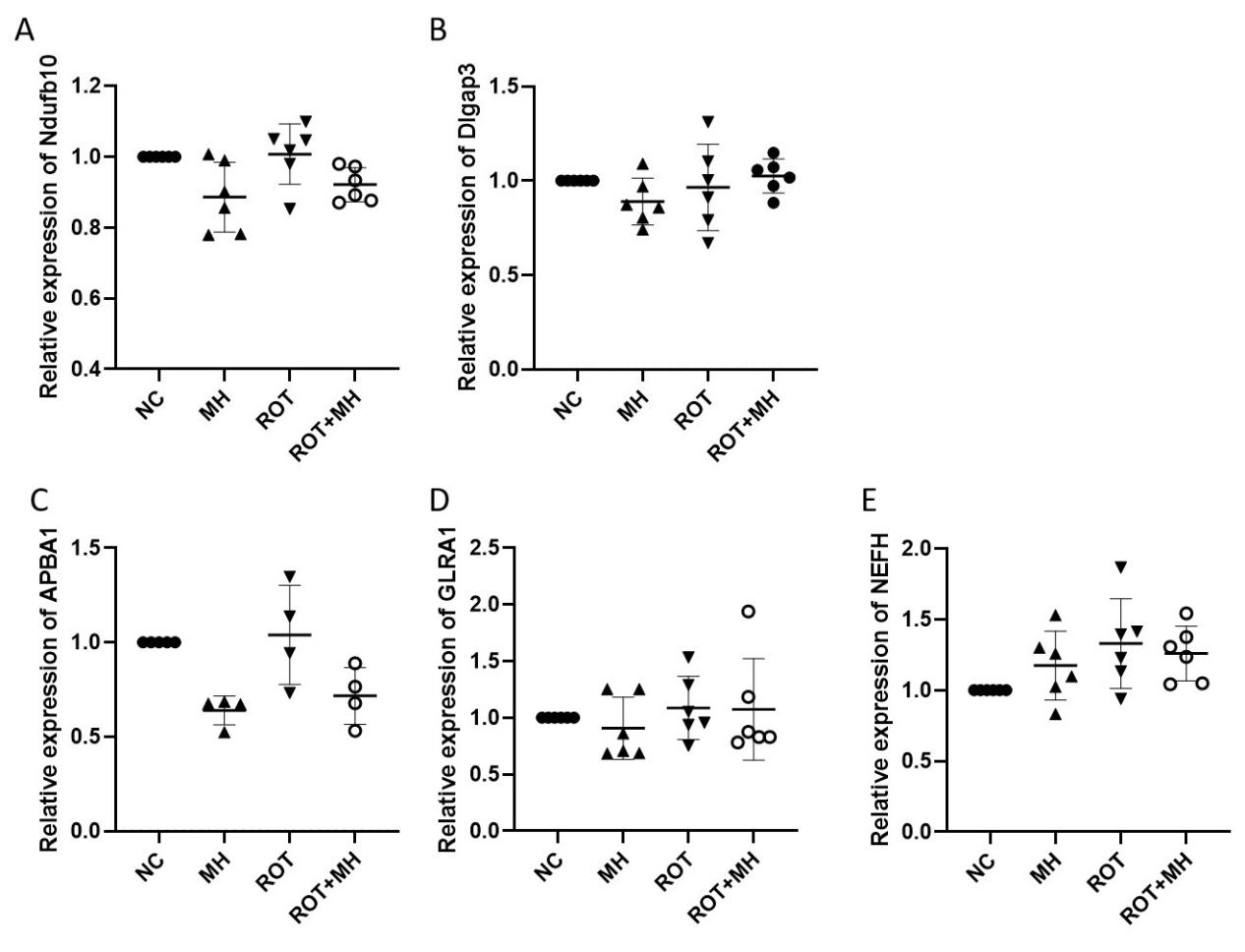
**Fig. S8. The effect of MH on regulating *Ndufb10, Dlgap3, APBA1, GLRA1* and *NEFH*.**
